# Cell signaling facilitates apical constriction by basolaterally recruiting Arp2/3 via Rac and WAVE

**DOI:** 10.1101/2024.09.23.614059

**Authors:** Pu Zhang, Taylor N. Medwig-Kinney, Eleanor A. Breiner, Jadyn M. Perez, April N. Song, Bob Goldstein

## Abstract

Apical constriction is a critical cell shape change that bends tissues. How precisely-localized actomyosin regulators drive apical constriction remains poorly understood. *C. elegans* gastrulation provides a valuable model to address this question. The Arp2/3 complex is essential in *C. elegans* gastrulation. To understand how Arp2/3 is locally regulated, we imaged embryos with endogenously-tagged Arp2/3 and its nucleation-promoting factors (NPFs). The three NPFs – WAVE, WASP, and WASH – colocalized with Arp2/3 and controlled Arp2/3 localization at distinct subcellular locations. We exploited this finding to study distinct populations of Arp2/3 and found that only WAVE depletion caused penetrant gastrulation defects. WAVE localized basolaterally with Arp2/3 at cell-cell contacts, dependent on CED-10/Rac. Establishing ectopic cell contacts recruited WAVE and Arp2/3, identifying contact as a symmetry-breaking cue for localization of these proteins. These results suggest that cell-cell signaling via Rac activates WAVE and Arp2/3 basolaterally, and that basolateral Arp2/3 is important for apical constriction.

## Introduction

Cell shape changes are fundamental to shaping animals during development. One of the most common cell shape changes is apical constriction – the shrinkage of the apical surface of a cell (Martin and Goldstein 2014). Apical constriction promotes tissue remodeling in various homeostatic and developmental contexts, such as gastrulation in many organisms (Martin and Goldstein 2014). Neural tube formation in vertebrates also depends on apical constriction, and defective neural tube formation is one of the most common classes of human birth defects (Nikolopoulou et al. 2017). Understanding the mechanisms underlying apical constriction is essential to understanding normal development and disease states.

Apical constriction is driven by the contraction of cortical actomyosin networks and the linkage of these networks to adherens junctions at the plasma membrane (Martin and Goldstein 2014; Clarke and Martin 2021). While there is some understanding of how spatially localized actomyosin regulators drive cytoskeletal rearrangement during processes like cell crawling and division, we know far less about how actomyosin networks are regulated during apical constriction. We predict that various actin regulators may have important roles in various sites in apically constricting cells, much as actin regulators do in crawling cells (Blanchoin et al. 2014). To address this knowledge gap, here we focused on the local regulation of an important actin regulator, the Arp2/3 complex, using *C. elegans* gastrulation as our model.

Gastrulation in *C. elegans* begins when the embryo is at the 26-cell stage. Two endodermal precursor cells (EPCs), called E anterior and E posterior (Ea/p), internalize from the surface of the embryo by apical constriction (Figure 1A) (Nance, Munro, and Priess 2003; Lee and Goldstein 2003).

**Figure 1.**
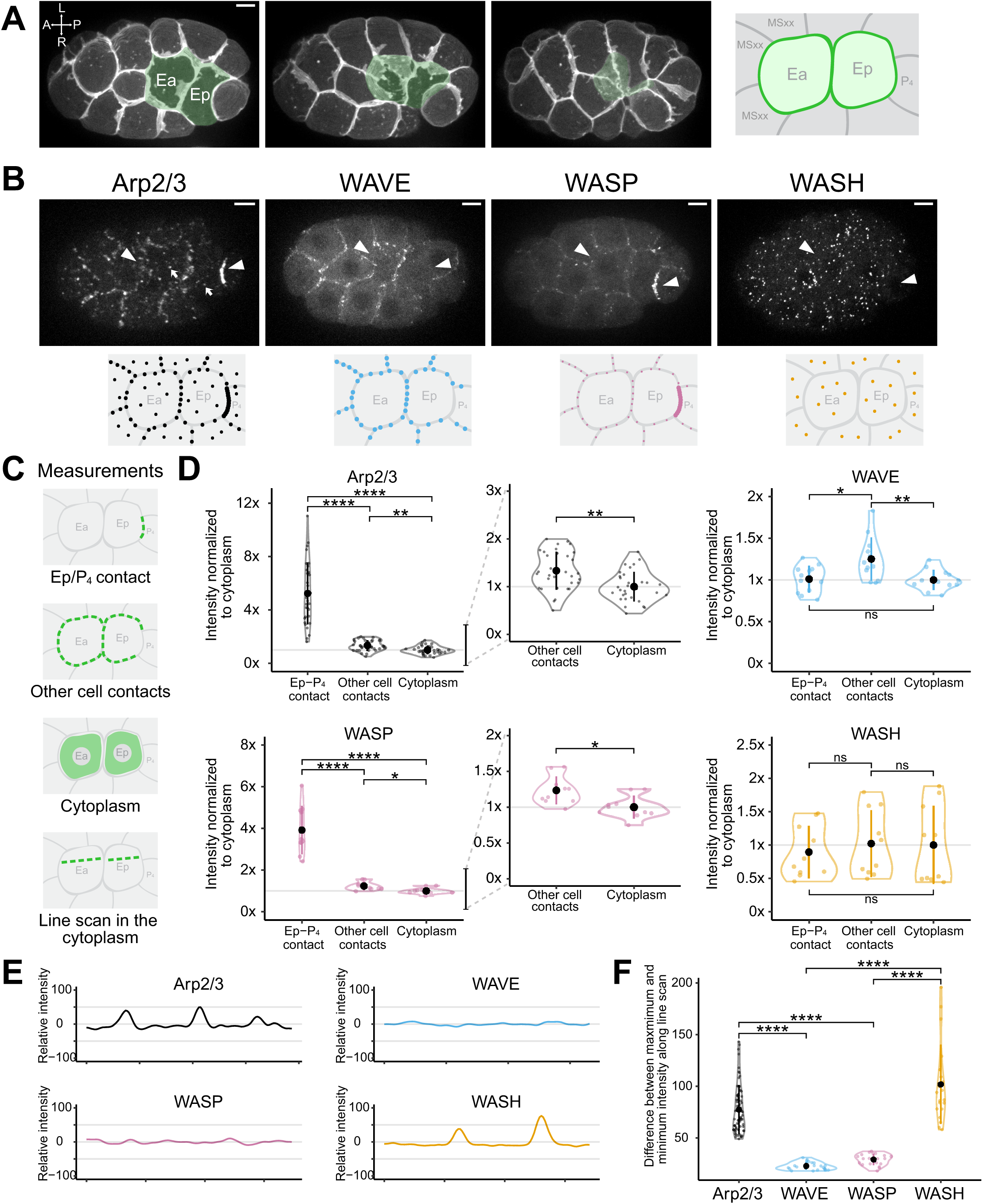
Quantitative analysis of Arp2/3 and nucleation-promoting factors (NPFs) localization during apical constriction. (A) Maximum intensity projections of 10 planes spanning a total Z-depth of 5 μm, depicting *C. elegans* gastrulation from a ventral view with plasma membranes fluorescently labeled (mex-5p::mScarlet-I::PH). Ea and Ep cells are pseudo-colored to visualize their internalization over time. The diagram to the right shows Ea and Ep, along with their neighboring cells, is used throughout the paper to depict cellular and subcellular protein localization. (B) Micrographs from time-lapse movies depicting localization of Arp2/3 (ARX-2::TagRFP), WAVE (GFP-C1^3xFlag::GEX-3), WASP (GFP::WSP-1A), and WASH (WSHC-5::mNG-C1^3xFlag) from a ventral view. White arrowheads point to Ea and Ep cells. White arrows point to vesicle-like structures in the cytoplasm. The diagrams underneath each micrograph highlight the observed localization in Ea, Ep, and neighboring cells. (C) Diagrams representing the four regions of interest for quantification: the Ep-P_4_ contact, the cell-cell contacts, the cytoplasm, and a line scan within the cytoplasm. (D) Violin plots depicting normalized fluorescence intensity of Arp2/3, WAVE, WASP, and WASH at Ep/P_4_ contacts, cell-cell contacts, and the cytoplasm. Insets of Arp2/3 and WASP highlight the difference between the signal at the cell-cell contacts and the cytoplasm. Measurements were collected 6 minutes after the division of neighboring mesoderm precursor cells (MSx). (center dot, mean; vertical line, standard deviation (s.d.); outline, the distribution of the data; n ≥ 10 embryos) (E) Representative line scan measurements of Arp2/3, WAVE, WASP, and WASH. (F) Violin plots depicting the differences between the maximum and minimum gray values along the line scan. (center dot, mean; vertical line, s.d.; outline, the distribution of the data; n ≥ 10 embryos) P-values reported in Figures D and F were calculated using either Tukey’s test or Dunnett’s test, depending on whether the variances between groups were equal. (*p<0.05, **p<0.01, ***p<0.001, ****p<0.0001) Scale bar: 5 µm.

Loss-of-function studies in diverse organisms have revealed the importance of the Arp2/3 complex in various morphogenetic events, including apical constriction in *C. elegans* – 100% of Arp2/3-depleted embryos form membrane blebs and exhibit a gastrulation-defective (Gad) phenotype, with EPCs failing to become fully covered by surrounding cells before the EPCs divide (Severson, Baillie, and Bowerman 2002; Roh-Johnson and Goldstein 2009). The Arp2/3 complex is an actin nucleator that binds existing actin filaments and initiates the formation of new filaments at a characteristic ∼70° angle (Mullins, Heuser, and Pollard 1998; Svitkina and Borisy 1999). Arp2/3-generated branched actin networks are present in the cell cortex and near the surface of many membranous organelles (Svitkina 2018). These networks generate pushing forces during processes such as cell migration and vesicular trafficking (Suraneni et al. 2012; Wu et al. 2012; Galletta, Chuang, and Cooper 2008).

Arp2/3 is activated by nucleation-promoting factors (NPFs) (Bieling and Rottner 2023). Moreover, different NPFs fulfill specialized functions at distinct subcellular locations (Molinie and Gautreau 2018). For instance, the WAVE complex at the leading edge of lamellipodia facilitates cell migration, particularly persistent directional migration (Steffen et al. 2004). N-WASP plays a crucial role in endocytosis at the plasma membrane (Merrifield et al. 2004; Benesch et al. 2005). The WASH complex activates Arp2/3 on the surface of endosomes, regulating endosomal sorting and trafficking (Gomez and Billadeau 2009; MacDonald et al. 2018). We were curious whether this division of roles also exists in *C. elegans* gastrulation, and if so, whether we could take advantage of this to further dissect which subcellular population of Arp2/3 is relevant for apical constriction, and how this population is locally regulated.

Whereas vertebrates typically have several NPF complexes, *C. elegans* has just three known NPF complexes – a WAVE complex, a WASP complex, and a WASH complex, with each complex member encoded by a single gene (Figure S1A) (Kollmar, Lbik, and Enge 2012; Smolyn 2020). This provided us with an opportunity to examine Arp2/3 regulation in apical constriction by studying a complete set of NPFs and avoid potential function redundancy between different isoforms (Tang et al. 2020). To answer how NPFs might control Arp2/3 activity in apical constriction, we visualized the localization of Arp2/3 along with WAVE, WASP, and WASH via live-cell imaging of tagged endogenous proteins. We observed that WAVE, WASP, and WASH had distinct subcellular locations. Further, these NPFs colocalized with Arp2/3 and controlled Arp2/3 localization at each of these subcellular locations. We then used the NPFs as tools to study the contributions of different populations of Arp2/3 in apical constriction. We confirmed that WAVE is required for gastrulation (Sullivan-Brown et al., 2016) and found that WASP makes a minor role that is fully redundant with WAVE, whereas depleting WASH had no detectable effect on gastrulation. To determine which signaling pathway(s) might activate WAVE and Arp2/3 at a subset of cell-cell contacts, we targeted Rac genes expressed in the early embryo since Rac small GTPases bind and activate WAVE in diverse systems (Rottner, Stradal, and Chen 2021). We identified a Rac that is required for WAVE and Arp2/3 localization to cell-cell contacts. These results lead to the hypothesis that cell-cell contacts serve as a symmetry-breaking cue that localizes WAVE and WAVE-activated Arp2/3 to sites on the basolateral but not apical cell cortex. We confirmed this model by creating ectopic cell-cell contacts and finding that the ectopic contacts were sufficient to recruit WAVE and Arp2/3. Our results suggest that cell-cell signaling via Rac and WAVE positions Arp2/3 basolaterally, where it makes an important contribution to apical constriction by mechanisms that are not yet understood.

## Results

### The Arp2/3 complex localizes to cell-cell contacts and vesicle-like structures in the cytoplasm

To elucidate which subcellular population(s) of Arp2/3 contribute to apical constriction, we first examined its localization. Our lab previously generated an antibody against one of the Arp2/3 subunits, ARX-5, to examine Arp2/3 distribution in fixed samples. We observed Arp2/3 enriched near plasma membranes in 26 to 28-cell stage embryos (Roh-Johnson and Goldstein 2009). To examine Arp2/3 localization over time using live-cell imaging, we filmed embryos with ARX-2 tagged with TagRFP at its C-terminus (Wu et al. 2017). The ARX-2::TagRFP strain was fully viable (Table S2), despite ARX-2 being an essential protein (Severson, Baillie, and Bowerman 2002), indicating that C-terminally tagged ARX-2 is functional. With endogenously tagged ARX-2, we now show that the Arp2/3 complex enriched near specific cell-cell contacts as well as vesicle-like structures in the cytoplasm (Figure 1B, white arrows point to vesicle-like structures in the cytoplasm). We observed a brighter signal where the internalizing endodermal precursor Ep and the germline precursor P_4_ contact each other, compared to other cell-cell contacts. To quantitatively analyze the distribution of ARX-2 over time, we collected time-lapse movies of multiple embryos and temporally aligned the movies using the birth of the neighboring MSxx cells (i.e. the four granddaughters of the MS cell) as a reference point. EPCs complete apical constriction approximately 18 minutes post-MSxx division, so we quantified the ARX-2 signal intensities at regular intervals before then: at 0, 6, and 12 minutes. We focused on three regions in Ea/p and their neighboring cells: the Ep-P_4_ contact, the contacts between Ea and Ep with each other and their neighboring cells excluding P_4_, and the cytoplasm (Figure 1C). We found that at all the time points analyzed, ARX-2 was most enriched at Ep/P_4_ contacts, with a signal intensity ∼5 times that of the cytoplasmic signal; ARX-2 was also enriched at other cell-cell contacts, although to a lesser extent: ARX-2 signal showed a punctate distribution at the non-Ep-P_4_ cell-cell contacts, and the average signal intensity along these cell-cell contacts, which included the punctae, was ∼1.3 times that of the cytoplasmic signal (Figure 1D and Figure S1B). Because the overall localization patterns were similar at all time points, we showed 6 minutes as a representative time point in Figure 1D and the rest of the time points were included in Figure S1B.

### Three NPFs exhibit distinct cellular and subcellular localization during apical constriction

Next, we aimed to determine how the Arp2/3 complex became localized at the aforementioned locations. Because different NPFs have been shown to fulfill specialized functions at distinct subcellular locations in migrating cells (Molinie and Gautreau 2018), we hypothesized that NPFs play critical roles during apical constriction by modulating Arp2/3 activities at specific sites. To test this hypothesis, we examined the localization of the three known NPFs in *C. elegans* – the WAVE, WASP, and WASH complexes – in early embryos. Specifically, we visualized tagged endogenous components of these complexes (Figure S1A). Using the same quantification pipeline as we used for the Arp2/3 complex, we analyzed the distribution of each NPF.

To examine the localization of the WAVE complex, we filmed embryos expressing GEX-3 tagged with GFP at its N terminus (Heppert et al. 2016). The tagged strain had an embryo hatching rate of 99%, indicating that the tagged GEX-3 was functional (Table S2). For simplicity, we refer to the *gex-3* gene and GEX-3 protein as WAVE hereafter. WAVE localized to most cell-cell contacts with an average signal intensity along the contacts approximately 1.25 times that of the cytoplasmic signal, except at the Ep/P_4_ contact, where the signal was indistinguishable from the cytoplasmic signal (Figure 1B, Figure 1D, and Figure S1B). This localization to cell-cell contacts during gastrulation is consistent with WAVE’s basolateral localization at later stages in epithelial cells of the epidermis (Patel et al. 2008; Bernadskaya et al. 2012).

The *wsp-1* locus is predicted to encode two protein isoforms, WSP-1A and WSP-1B, using two different transcriptional start sites. We used a strain expressing WSP-1A tagged with GFP at its N terminus (Wu et al. 2017) and tagged WSP-1B with mNeonGreen at its N terminus using CRISPR/Cas9-dependent genome editing. Both strains were fully viable (Table S2). WSP-1A primarily localized to the Ep/P_4_ contact, with a signal intensity about 4 times that of the cytoplasmic signal. WSP-1A is also enriched at other cell-cell contacts, with an average signal intensity along contacts approximately 1.25 times that of the cytoplasmic signal (Figure 1B, Figure 1D, and Figure S1B). WSP-1B was at levels below the detectable threshold on our imaging system, appearing similar to unlabeled, wild-type N2 control embryos at the time of Ea/Ep internalization (Figure S1C-E). For simplicity, we refer to the *wsp-1* locus and WSP-1A protein as WASP.

To examine the localization of the WASH complex, we used CRISPR/Cas9-dependent genome editing to tag *wshc-5* with mNeonGreen at its C terminus. The tagged strain is fully viable, indicating it was functional (Table S2). For simplicity, we refer to the *wshc-5* gene and WSHC-5 protein as WASH hereafter. WASH was not enriched at any cell-cell contacts but localized to vesicle-like structures in the cytoplasm (Figure 1B). Since WASH localization had not been documented previously in *C. elegans*, we compared our findings with observations from other systems. WASH is known to interact with F-actin capping proteins (Jia et al. 2010; Hernandez-Valladares et al. 2010), so we performed colocalization analyses between WASH and the capping protein CAP-1 with a dual-labeled strain (Figure S1F). Our analysis focused on the previously defined regions: the Ep-P_4_ contact, the other cell-cell contacts, and the cytoplasm (Figure 1C). We measured and compared the Pearson correlation coefficient (PCC) at each of these locations for WASH and CAP-1: WASH and CAP-1 colocalized most strongly at punctate structures in the cytoplasm. This is consistent with previous work in other systems that found WASH localized to endocytic vesicles (MacDonald et al. 2018).

We proceeded to quantify WASH localization. The total signal intensities at all measured locations showed no difference, likely because the contribution from the vesicle-like structures is too weak to significantly affect the total cytoplasmic intensity measurements (Figure 1D and Figure S1B). Therefore, to quantify the signal distribution within the cytoplasm for WASH and other proteins, we performed line scans across the cytoplasm of Ea and Ep (Figure 1C). Because the overall localization patterns were similar at all time points based on our previous results (Figure 1D and Figure S1B), we used 6 minutes as a representative time point for this and the rest of the quantification in this study. We found that the signals for both WASH and Arp2/3 showed peaks along the line scan corresponding to the sites of the vesicle-like structures, whereas WAVE and WASP exhibited a more uniform distribution (Figure 1E). We used the difference between the maximum and minimum intensity along the line scan to quantify the degree of punctate signal distribution in the cytoplasm. We found that WASH and Arp2/3 had a consistently larger intensity difference compared to WAVE and WASP (Figure 1F).

We conclude that three NPFs in *C. elegans* localize to mostly distinct subcellular locations, with some overlap between WAVE and WASP at cell-cell contacts.

### Three NPFs occupy distinct Arp2/3-enriched cellular and subcellular locations

To investigate whether the three NPFs colocalize with Arp2/3 at their respective subcellular locations, we performed live-cell imaging of dual-labeled strains for each NPF and Arp2/3 (Figure 2A). We conducted colocalization analyses using the quantification pipeline that we established for WASH and CAP-1. Each of the NPFs colocalized with Arp2/3. WAVE and Arp2/3 colocalized most strongly at the non-Ep-P_4_ cell-cell contacts (Figure 2B). WASP and Arp2/3 exhibited the strongest colocalization at the Ep-P_4_ contact, with a weaker but still significant colocalization at the other cell-cell contacts, and no obvious colocalization in the cytoplasm (Figure 2B). WASH and Arp2/3 colocalized most strongly in the cytoplasm (Figure 2B).

**Figure 2.**
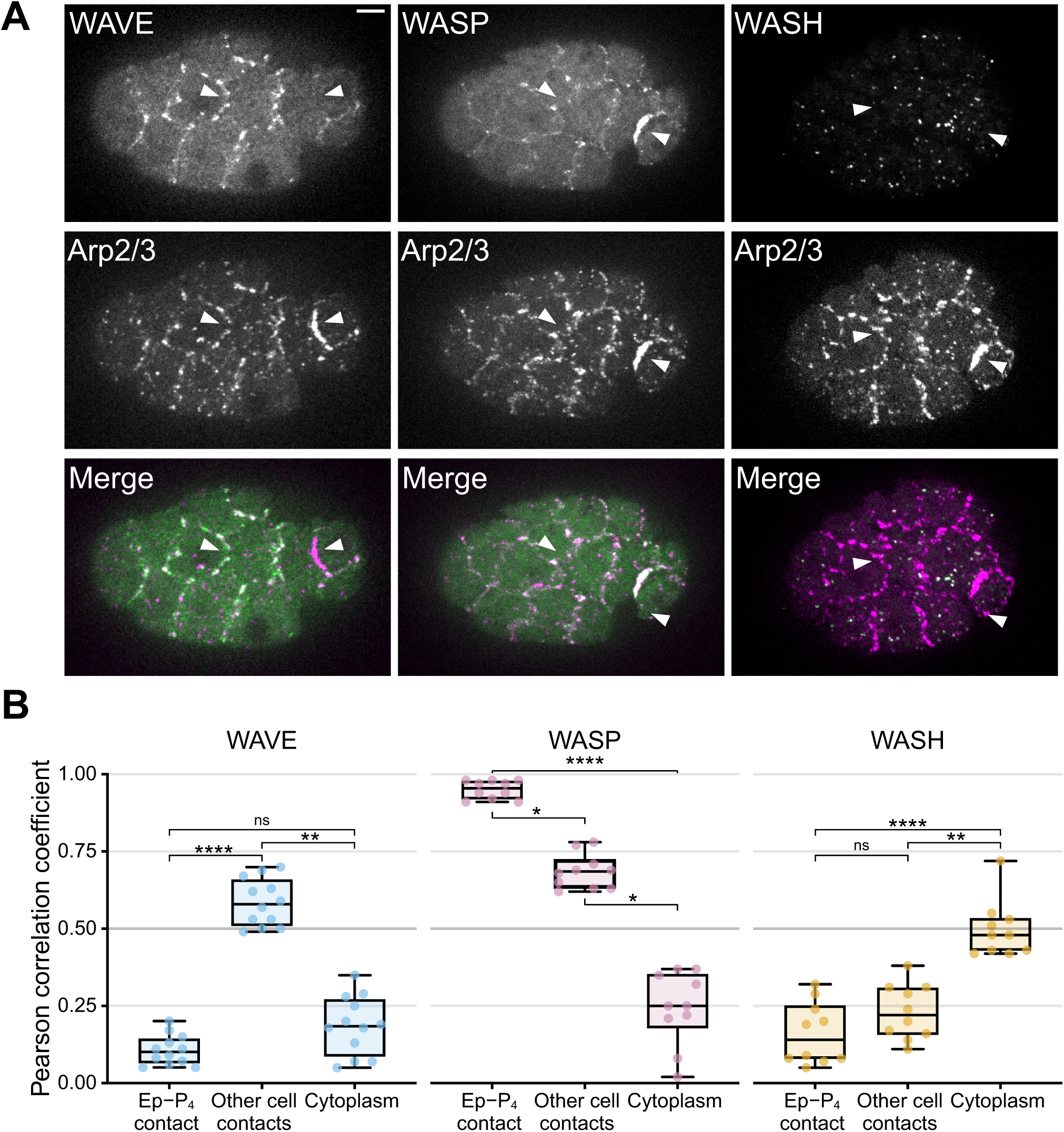
Three NPFs colocalize with Arp2/3 at different cellular and subcellular locations. (A) Co-visualization of Arp2/3 with WAVE, WASP, and WASH in gastrulation stage embryos using endogenously tagged alleles. White arrowheads point to Ea and Ep cells. Scale bar: 5 µm. (B) Quantification of colocalization between Arp2/3 and NPFs, with box plots reporting Pearson correlation coefficients at Ep/P_4_ contacts, cell-cell contacts, and the cytoplasm. P values were calculated with a Dunn’s test. (center line, median; box, interquartile range (IQR); whiskers, min/max range; n ≥ 10 embryos; *p<0.05, **p<0.01, ***p<0.001, ****p<0.0001)

### NPFs build Arp2/3-enriched structures at different cellular and subcellular locations

Our results so far led us to hypothesize that NPFs modulate Arp2/3 activities at distinct subcellular locations, with potential functional overlap between WAVE and WASP at cell-cell contacts. To test this hypothesis, we observed and quantified Arp2/3 localization after depleting the NPFs by RNAi. We maximized RNAi efficiency by synthesizing double-stranded RNAs using *C. elegans* cDNA as templates and performing RNAi by injection. Then, we quantitatively assessed the extent of protein knockdown by imaging and quantifying the total fluorescence from three sets of embryos side by side: unlabeled (wild-type) embryos, embryos expressing the fluorescent tags, and embryos with tagged components and targeted by dsRNA. In all cases, the total fluorescence in the tagged, knockdown embryos was indistinguishable from that in unlabeled embryos, indicating that RNAi was effective at depleting these proteins to undetectable levels (Figure S2A, B).

We then depleted each NPF in Arp2/3-labeled embryos and quantified the change in Arp2/3 localization by imaging and comparison of RNAi-treated and control embryos side by side. In Figures 1 and 2, we imaged embryos from an en-face ventral view to capture the details of protein localization. While lateral view imaging provides less detail overall, it offers better visualization of protein distribution along the apical-basal axis. Since our results suggest that the NPFs and Arp2/3 localize to cell-cell contacts and within the cytoplasm, we switched to the lateral view for subsequent imaging (Figure 3A). WAVE(RNAi) reduced the Arp2/3 signal at the non-Ep/P_4_ cell-cell contacts to the cytoplasmic level while leaving the signal at the Ep-P_4_ contact and in the cytoplasm mostly unchanged (Figure 3A, B). WASP(RNAi) reduced the Arp2/3 signal at the Ep/P_4_ contact to the cytoplasmic level. Arp2/3 at the non-Ep/P_4_ cell-cell contacts was not affected by WASP(RNAi), and the signal in the cytoplasm increased, likely due to the failure to recruit Arp2/3 to the Ep-P_4_ contact (Figure 3A, B). WASH RNAi eliminated most of the cytoplasmic, vesicle-like Arp2/3 signal as observed through live imaging (Figure 3A). Our quantification method detected no changes at any measured locations (Figure 3B), consistent with our previous observations (Figure 1D and Figure S1B). Therefore, to quantify the effect of WASH(RNAi), we again implemented the line scan method to capture the change in signal intensity in the cytoplasm. WASH(RNAi) reduced the difference between maximum and minimum Arp2/3 signal intensity along the line scan, whereas WAVE(RNAi) and WASP(RNAi) led to no obvious change (Figure 3C).

**Figure 3.**
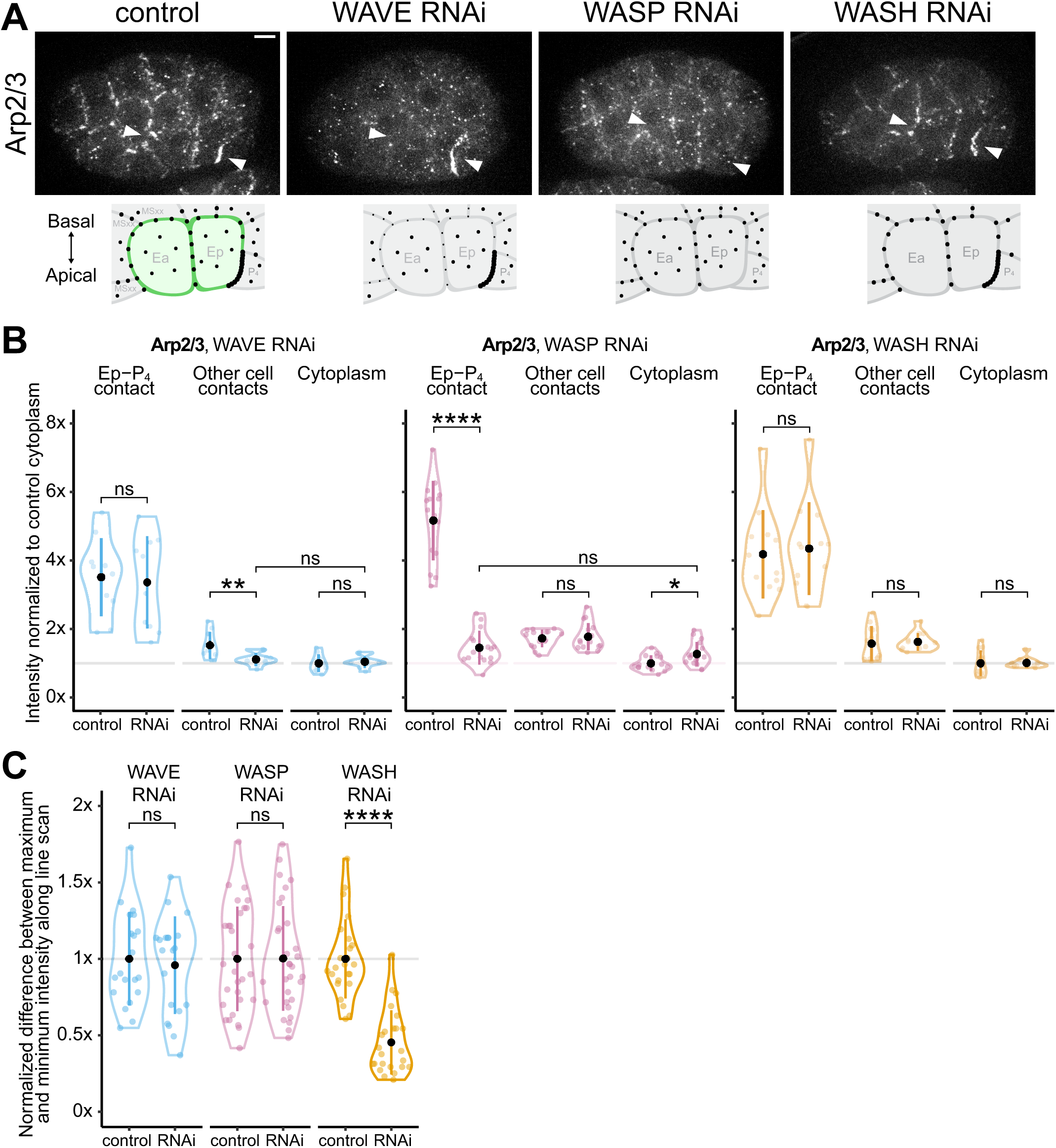
RNAi against each NPF leads to Arp2/3 reduction at different cellular and subcellular locations. (A) Micrographs from time-lapse movies depicting localization of Arp2/3 in control and WAVE, WASP, or WASH RNAi-treated embryos from a lateral view. White arrowheads point to Ea and Ep cells. The diagrams underneath each micrograph highlight the observed Arp2/3 localization in E and neighboring cells. (B) Violin plots reporting changes in Arp2/3 localization at Ep/P_4_ contacts, cell-cell contacts, and the cytoplasm upon RNAi depletion of WAVE, WASP, and WASH. (center dot, mean; vertical line, s.d.; outline, the distribution of the data; n ≥ 10 embryos) (C) Violin plots reporting changes in the differences between the maximum and minimum gray values of the Arp2/3 signal along the line scan upon RNAi depletion of WAVE, WASP, and WASH. (center dot, mean; vertical line, s.d.; outline, the distribution of the data; n ≥ 10 embryos) All measurements were collected 6 minutes after the division of neighboring mesoderm precursor cells (MSx). P-values reported in Figures B and C were calculated using either an unpaired t-test or an unpaired t-test with Welch’s correction, depending on whether the variances between groups were equal. (*p<0.05, **p<0.01, ***p<0.001, ****p<0.0001) Scale bar: 5 µm.

We conclude that discreet NPFs build Arp2/3-enriched structures at distinct cellular and subcellular locations: WAVE is mainly controlling Arp2/3 at the non-Ep/P_4_ cell-cell contacts, WASP is responsible for the signal at the Ep/P_4_ contact, and WASH is responsible for Arp2/3 at the vesicle-like structures.

### Depletion of each NPF and depleting multiple NPFs in combination led to different degrees of gastrulation defects

Our RNAi results positioned us to study the contributions of different subcellular populations of Arp2/3 in apical constriction. We investigated how NPF deletion might affect *C. elegans* gastrulation using RNAi against each NPF alone or in combination. Embryos from injected worms were filmed using Differential Interference Contrast (DIC) microscopy and examined for gastrulation defects, defined as the failure of the two EPCs to fully internalize, leaving part of at least one EPC uncovered by the neighboring cells before division.

Consistent with our previous findings (Sullivan-Brown et al. 2016), WAVE RNAi alone resulted in a highly penetrant gastrulation-defective phenotype, in 40 out of 46 (87%) embryos (Figure 4A, B). Depleting WASP and WASH individually or in combination did not disrupt gastrulation (Figure 4A, B). However, combining WASP(RNAi) with WAVE(RNAi) significantly enhanced the WAVE(RNAi) phenotype, with 63 out of 64 (94%) embryos showing gastrulation defects (Figure 4A, B), indicating that WASP is fully redundant with WAVE in apical constriction. Deleting all NPFs simultaneously resulted in 100% gastrulation defects in all 44 embryos examined, mirroring the 100% defect observed in Arp2/3 RNAi embryos (Figure 4A, B). Since the premature division of the EPCs can prevent their internalization (Lee et al. 2006), we examined the timing of Ea/p division by measuring the time between the MS division and the Ea/p division (Table S4). None of the RNAi treatments led to a significant change in division timing, indicating that Ea and Ep did not divide prematurely in NPF-depleted embryos. Taken together, our results suggest that the subpopulation of Arp2/3 activated by WAVE at the non-Ep/P_4_ cell-cell contacts is the main contributor to apical constriction (Figure 4B). We next aimed to understand how WAVE becomes activated at these cell-cell contacts.

**Figure 4.**
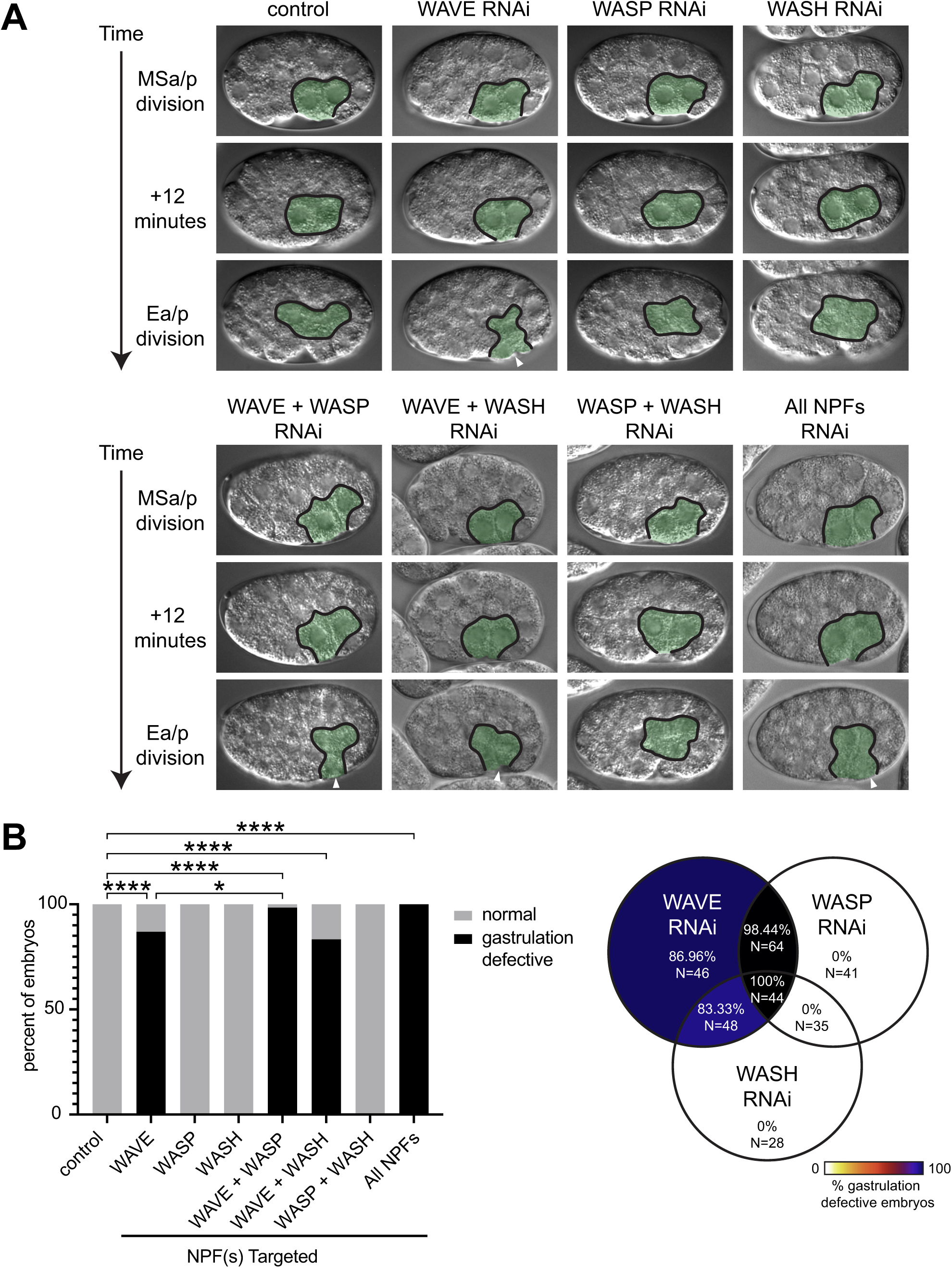
NPF RNAi by themselves and in combination lead to different degrees of gastrulation defects. (A) Micrographs from time-lapse DIC movies in eight different backgrounds, with time indicated on the left from the MSx cell division. E lineage cells are outlined and pseudocolored in green. Gastrulation defects (E cells dividing before being fully covered by neighboring cells) are indicated with white arrowheads. An enclosed outline and the absence of arrowheads indicate that endodermal precursors became internalized at the 2E stage, as seen in wild-type embryos. (B) The bar graph (left) and Venn diagram (right) summarize the effects of NPF RNAi, both individually and in combination, on gastrulation. The heat map represents the different penetrance levels, with darker colors indicating a higher percentage of gastrulation defects. P values were calculated using Fisher’s exact test. (**\*p<0.05**, **p<0.01, ***p<0.001, ****p<0.0001)

### WAVE is activated by a Rac1 GTPase, CED-10, at cell-cell contact sites

In migrating cells and epithelial cells, WAVE is known to be activated by Rac family GTPases at the leading edge (Miki, Suetsugu, and Takenawa 1998; Innocenti et al. 2004; Bernadskaya et al. 2012). To determine if Rac also regulates WAVE during apical constriction, we first checked the expression of Rac genes in early embryos and the germline using published RNAseq databases from our lab and others. We found that two of the three putative Rac genes, *ced-10* and *mig-2*, are expressed in early embryos and the germline, whereas *rac-2* showed no detectable expression at these stages (Tintori et al. 2016; Diag et al. 2018). We then RNAi-depleted *ced-10* or *mig-2* in WAVE-labeled embryos and quantified WAVE localization. Depletion of CED-10 reduced the WAVE signal at cell-cell contacts to the level of cytplasmic WAVE (Figure 5A, B). *mig-2(RNAi*) did not alter WAVE localization (Figure S3A, B).

**Figure 5.**
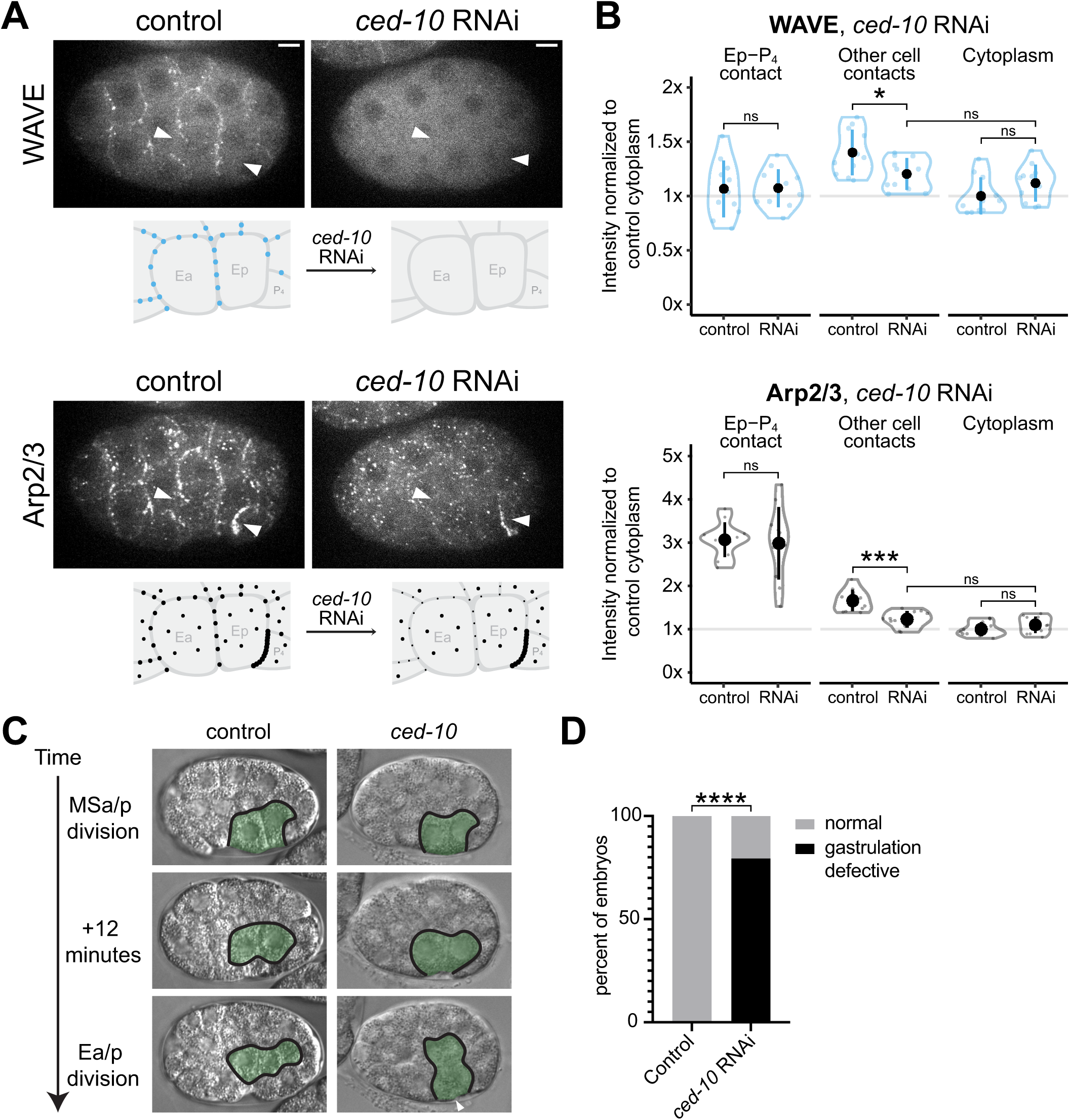
The Rac1 GTPase CED-10 recruits WAVE and Arp2/3 at cell-cell contact and contributes to apical constriction. (A) Micrographs from time-lapse movies depicting localization of WAVE and Arp2/3 in control and CED-10 RNAi-treated embryos from a lateral view. Scale bar: 5 µm. The diagrams underneath each micrograph highlight the observed Arp2/3 localization in E and neighboring cells. (B) Violin plots reporting changes in WAVE and Arp2/3 localization at Ep/P_4_ contacts, cell-cell contacts, and the cytoplasm upon RNAi depletion of CED-10. (center dot, mean; vertical line, s.d.; outline, the distribution of the data; n = 11 embryos) (C) Micrographs from time-lapse DIC movies of wild-type and CED-10 RNAi-treated embryos with time on the left from MSa/p cell division. E lineage cells are outlined and pseudocolored in green. Gastrulation defects (E cells dividing before being fully covered by neighboring cells) are indicated with white arrowheads. (D) Bar graph summarizing gastrulation defects caused by ced-10 RNAi. P-values reported in Figures B were calculated using either an unpaired t-test or an unpaired t-test with Welch’s correction, depending on whether the variances between groups were equal. P values reported in Figure D were calculated with Fisher’s exact test. (*p<0.05, **p<0.01, ***p<0.001, ****p<0.0001)

These findings indicate that CED-10 is crucial for recruiting WAVE to cell-cell contacts in the early embryo. We then explored if CED-10 controls Arp2/3 localization and found that *ced-10* RNAi reduced the Arp2/3 signal at cell-cell contacts to the cytoplasmic level, while the signal at Ep-P_4_ contact and in the cytoplasm remained mostly unchanged (Figure 5A, B). Since depletion of CED-10 or WAVE resulted in similar reductions in Arp2/3 levels (*i.e.*, down to cytoplasmic levels), we conclude that CED-10 is key in activating Arp2/3 at specific cell-cell contacts. To examine the localization of CED-10, we tagged it with mNeonGreen at the N terminus. This stable new strain has a hatching rate of 91% (Table S2). Because *ced-10 null* worms are not viable (Lundquist et al. 2001), we concluded that the tagged protein is partially functional. Live imaging of tagged endogenous CED-10 showed that it localizes to all cell-cell contacts, including the Ep-P_4_ contact (Figure S3C). Using this tagged endogenous CED-10, we verified that *ced-10*(*RNAi)* was highly effective at depleting CED-10 protein, to undetectable levels (Figure S3D-E).

Because of the role of CED-10 in recruiting WAVE and Arp2/3 to a subset of cell-cell contacts, we hypothesized that CED-10 function contributes to *C. elegans* gastrulation. To test this, we examined *ced-10* RNAi embryos by DIC imaging and found that 31 out of 39 targeted embryos exhibited gastrulation defects (Figure 5C, D). Taken together, our results demonstrate that the CED-10/Rac GTPase recruits WAVE and Arp2/3 to specific cell-cell contacts and plays a crucial role in gastrulation.

### Cell-cell contact acts as a symmetry-breaking cue to recruit WAVE and Arp2/3

Our results suggested that the Rac-WAVE-Arp2/3 signaling axis is activated at a subpopulation of cell-cell contacts. We hypothesized that cell-cell contacts serve as a symmetry-breaking cue to localize WAVE and Arp2/3 specifically to basolateral sites. To test this, we removed the eggshell and envelope from developing early embryos and created new cell-cell contacts by placing pairs of embryos in contact (Figure 6A). Both WAVE and Arp2/3 localized to the ectopic contacts (Figure 6A). The signal intensity at the ectopic contacts was significantly higher than the normally contact-free cell apex and was comparable to the endogenous contacts (Figure 6B). We conclude that cell-cell contacts are sufficient to recruit WAVE and Arp2/3 to sites of cell-cell contact. Because cell contacts normally define basolateral sites and we could recruit WAVE and Arp2/3 to previously contact-free apical sites by ectopically providing cell contacts there, we conclude that cell-cell contact acts as an apico-basal symmetry breaking cue that recruits WAVE and Arp2/3 specifically to basolateral sites.

**Figure 6.**
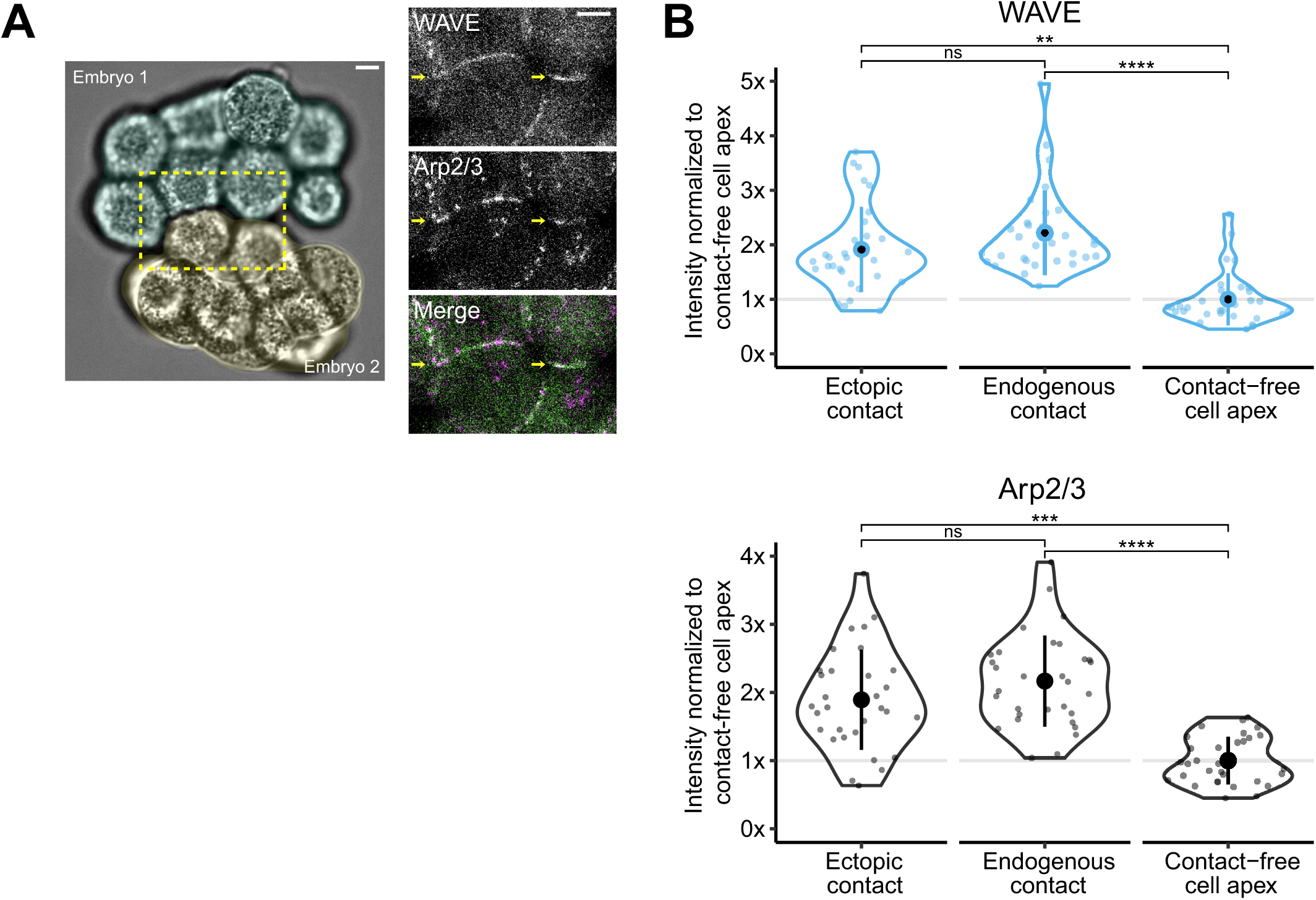
Newly established ectopic cell-cell contact acts as a symmetry-breaking cue to recruit WAVE and Arp2/3. (A) Micrographs from time-lapse movies of chimeric embryos created by combining two embryos expressing WAVE and Arp2/3. The DIC channel of the whole chimera is shown on the left, and two fluorescence channels of a blow-up of the chimeric contact (outlined region) are shown on the right. Yellow arrows point to ectopic cell-cell contacts. (B) Violin plots reporting normalized fluorescence intensity of WAVE and Arp2/3 at ectopic/endogenous cell-cell contacts and the contact-free cell apex. (center dot, mean; vertical line, s.d.; outline, the distribution of the data; n = 15 chimeras) P values were calculated with a Tukey test. (*p<0.05, **p<0.01, ***p<0.001, ****p<0.0001) Scale bar: 5 µm.

## Discussion

The actin cytoskeleton is crucial for generating and transmitting forces to neighboring cells in apical constriction (Lee and Goldstein 2003; Martin, Kaschube, and Wieschaus 2009; Slabodnick et al. 2023). However, we still have a limited understanding of the degree to which actin regulation in various subcellular locations contributes to apical constriction. Here we focused on one of the major nucleators of the actin cytoskeleton, the Arp2/3 complex, to gain insights into its function and regulation in *C. elegans* gastrulation. We discovered that three nucleation-promoting factors, the WAVE, WASP, and WASH complexes, activate distinct subpopulations of Arp2/3 at specific cellular and subcellular locations. WAVE RNAi, but not WASP or WASH RNAi, recapitulated the gastrulation defect caused by Arp2/3 RNAi, suggesting that the Arp2/3 complex activated by WAVE at a subset of cell-cell contacts is key in apical constriction. We identified that a Rac protein, CED-10, is responsible for localizing WAVE and Arp2/3 at these cell-cell contacts. Further, we found that ectopic cell-cell contacts between embryos could recruit WAVE and Arp2/3, revealing cell-cell contacts as an apico-basal symmetry breaking cue for localization of these proteins. In summary, our work reveals that cell-cell signaling makes an essential contribution to apical constriction by basolaterally recruiting Arp2/3 via Rac and WAVE, as depicted in the model in Figure 7A-B.

**Figure 7.**
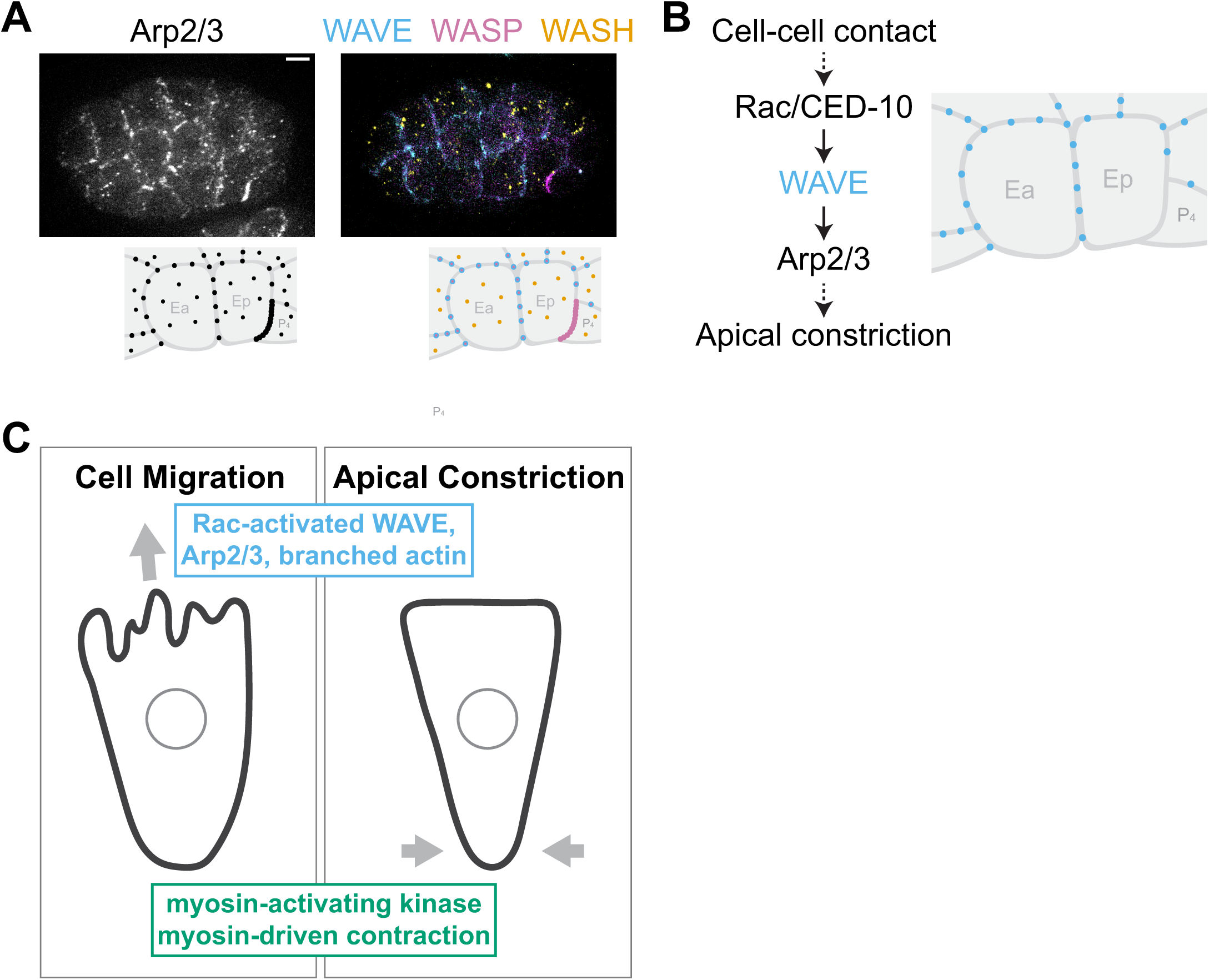
Summary. (A) The three NPFs in *C. elegans*, WAVE, WASP and WASH complexes, colocalize with Arp2/3 and control Arp2/3 localization at distinct subcellular locations. (B) Cell-cell signaling makes an essential contribution to apical constriction by basolaterally recruiting Arp2/3 via Rac and WAVE. (C) Rac signaling and myosin-activating kinase are active at opposing locations in migrating and apically constricting cells.

Components of the WAVE complex including Nckap1 and Abi1, as well as Rac1, have been implicated in neural tube closure in mice, with disruptions in these genes leading to neural tube defects (Rakeman and Anderson 2006; Migeotte, Grego-Bessa, and Anderson 2011; Dubielecka et al. 2011). Although the mechanisms behind these defects are not yet fully understood, our findings indicate that *C. elegans* gastrulation is a valuable model for further investigating underlying molecular and cell biological mechanisms.

Although our work reveals a subcellular population of Arp2/3 that contributes to gastrulation and reveals cellular and molecular mechanisms by which this specific population is specified, we do not yet understand how basolateral Arp2/3 contributes mechanistically to apical constriction. Arp2/3-mediated actin polymerization and myosin II-dependent contractility have been shown to play opposing roles in regulating cortical tension: active myosin II at the cortex slides actin filaments with respect to one another and generates cortical tension, whereas activation of the Rac-WAVE-Arp2/3 signaling axis decreases cortical tension (Tinevez et al. 2009; Bergert et al. 2012; Chugh et al. 2017). Elevated myosin II activity and cortical tension can lead to bleb formation, characterized by local detachment or rupture of the cortex from the plasma membrane (Tinevez et al. 2009). CED-10, WAVE, and Arp2/3-depleted embryos form membrane blebs, which may indicate elevated membrane tension and/or decreased membrane-cortex linkage (Roh-Johnson and Goldstein 2009; Sullivan-Brown et al. 2016; Lamb et al. 2024). To test if the blebbing phenotype is linked to the gastrulation defects seen in WAVE-depleted embryos, we attempted to rescue the gastrulation defects by tuning myosin levels with partial RNAi. Using a strong loss-of-function strain of GEX-3/WAVE, WM43, we found that reducing myosin levels with partial RNAi could restore membrane morphology and partially rescue gastrulation defects in WAVE-depleted embryos (Figure S4A, B), suggesting to us that the gastrulation defects caused by WAVE depletion may be related to the blebbing phenotype and membrane-cortex linkage issues. We speculate that active Rac-WAVE-Arp2/3 signaling at cell-cell contacts is crucial for maintaining proper membrane-cortex linkage, and that strong linkage at sites of cell-cell contact is important for successful apical constriction.

Our genetic analysis indicated that WASP makes a minor, redundant contribution to apical constriction. This may reflect the importance of strong Arp2/3 enrichment at the Ep/P_4_ contact. Alternatively, WASP might recruit a small amount of Arp2/3 at other cell-cell contacts, an amount that was below the detection limit of our current quantification methods. A functional coordination between WAVE and WASP has been shown in C. elegans neuroblast migration – WAVE mutations impair migration, while WASP mutations alone do not affect migration but exacerbate defects in WAVE-deficient cells (Zhu et al. 2016). Our findings indicate that WAVE and WASP may be coordinating with each other at a cellular and/or subcellular level in apical constriction. In the future, we are interested in understanding the striking enrichment of WASP and the exclusion of WAVE at the Ep/P_4_ contact and its potential role in apical constriction and other processes, such as asymmetric division. Additionally, this may also help explain the previously observed enrichment of capping protein at the endoderm/germline precursor cells’ contacts (Zhang, Medwig-Kinney, and Goldstein 2023; Goldstein 2000).

WASH depletion did not result in gastrulation defects. However, it has been shown to play a role in *C. elegans* embryonic development, perhaps by affecting processes that happen at later stages (Smolyn 2020). We found that WASH colocalizes with the capping protein CAP-1, as shown in other systems. CAP-1 has been observed surrounding the germ cell nuclei (Ray et al. 2023), and the authors of this paper speculated that this perinuclear localization is due to the capping protein being a component of the dynactin complex. Our findings suggest an alternative explanation for this localization: CAP-1 may localize to these sites as a member of the WASH complex. These possibilities are not mutually exclusive: WASH has been reported to assemble a super complex with dynactin where the Capping Protein (CP) is exchanged from dynactin to the WASH complex (Fokin and Gautreau 2021). The distinct perinuclear location in early embryos may offer an opportunity to further dissect the function and organization of these complexes.

The quantification pipeline we developed in this paper will be valuable for analyzing and understanding the function and regulation of other actin regulators during apical constriction, particularly those active at similar cellular locations, including the capping proteins. The regulation and interplay between different NPFs play crucial roles in processes such as directional cell migration, membrane remodeling, and vesicular trafficking. Our results highlight their distinct contributions to apical constriction by regulating the Arp2/3 complex. The system we present here offers a valuable tool for scientists interested in NPF-dependent processes to dissect their regulation and activation, potentially revealing organizing principles beyond the context of apical constriction.

The Rac-WAVE-Arp2/3 signaling axis plays crucial roles in various fundamental processes, including lamellipodia formation, junction establishment and maintenance, and regulation of cortical tension (Steffen et al. 2004; Yamazaki, Oikawa, and Takenawa 2007; Bergert et al. 2012; Sasidharan et al. 2018). In migrating cells, Rac activates Arp2/3 at the leading edge to produce branched-actin networks and generate pushing force. Myosin II, on the other hand, typically gets activated at the rear end of migrating cells and contributes to trailing edge retraction and steering the direction of migration (Ridley et al. 2003; Vicente-Manzanares et al. 2007; Allen et al. 2020). Intriguingly, in apically constricting cells in *C. elegans* embryos, Rac signaling and a myosin-activating kinase are also active at distinct locations: WAVE and Arp2/3 localize basolaterally via cell-cell contacts, while myosin is enriched at the contact-free apices of Ea and Ep cells (Figure 7C). Because lamellipodia-based cell crawling is found more broadly than in metazoans and therefore is likely to have evolved earlier than apical constriction (Brunet 2023), the similarity between the two systems could feasibly reflect an ancient evolutionary co-opting of cell crawling mechanisms for bending cell sheets. Consistent with this possibility, Rac and Rho function antagonistically in both crawling cells and some apically constricting cells (Chauhan et al. 2011; Nobes and Hall 1995; Burridge and Wennerberg 2004). Alternatively, lamellipodia-based cell crawling and apical constriction might have evolved independently. Due to limitations of resolution and imaging depth, we were not able to resolve whether WAVE and Arp2/3 contribute to treadmilling structures extended from the basolateral sides of cells. In the future, newer techniques may clarify whether active migration at the basolateral side of cells contributes to apical constriction.

## Material and methods

### Strain maintenance

Table S1 lists the strains used in this study. Some strains were provided by the CGC, which is funded by NIH Office of Research Infrastructure Programs (P40 OD010440). Worms were cultured on Nematode Growth Medium plates at 20 to 22°C following standard protocols (Brenner 1974; Sternberg et al. 2024).

### CRISPR-mediated genome engineering

The new endogenously-tagged strains described in this work were created using two previously established CRISPR/Cas9-mediated genome editing protocols: a Cas9 protein-based method (Ghanta and Mello 2020) and a Cas9-expressing plasmid method (Dickinson et al. 2015). Cas9 targeting sequences for each gene were selected using the CRISPR design tool CRISPOR (Concordet and Haeussler 2018). In the protein-based method, tracrRNA and crRNA were mixed with Cas9 protein (all reagents from IDT) and incubated at 37°C for 15 minutes. Then, a double-stranded DNA donor, containing codon-optimized mTurquoise2 (amplified from pDD315, AddGene #73343), mNeonGreen (amplified from pDD268, AddGene #132523), or mScarlet-I (amplified from pMS050, AddGene #91826) with 35 bp homology arms, was melted, cooled, and added to the mixture. Additionally, pRF4 (*rol-6(su1006)*) plasmid was included as a coinjection marker. The injection mix was then centrifuged, transferred to a fresh tube, and kept on ice during the injection. Heterozygous mutants were selected by genotyping F1 non-rollers or examining F1 under a Zeiss Axiozoom V16 stereo microscope. In the plasmid-based method, repair templates were constructed by integrating homology arm PCR products, which were amplified from worm genomic DNA, into vectors containing a fluorescent protein and a selection cassette using Gibson Assembly (New England Biolabs), as thoroughly detailed by Dickinson et al. (Dickinson et al. 2015). Cas9 guide sequences were inserted into the Cas9-sgRNA expression vector pDD162 and co-injected into the adult germlines along with the repair template vector and array markers. The selection of the edited worms was performed using previously established techniques (Dickinson et al. 2015). Homozygotes made with both methods were sequenced to confirm the edits. sgRNAs and homology arms sequences can be found in Table S3-1.

### RNA interference

Primers were designed to amplify approximately 1000 bp of the protein-coding sequence of each target gene (Table S3-2). cDNA from wild-type N2 worms was prepared using SuperScript™ III First-Strand Synthesis SuperMix according to the manufacturer’s instructions (Invitrogen). Each primer included 15 bases of the T7 promoter sequence at the 5’ end for use in a 2-step PCR with the wild-type cDNA. The PCR product was purified using a Zymoclean Gel DNA Recovery Kit (Zymo Research) and used as a template for a second PCR with primers containing the full-length T7 promoter sequence. After another round of purification with the Zymoclean Gel DNA Recovery Kit, the PCR product served as a template in a T7 RiboMAX Express RNAi System (Promega) following the manufacturer’s instructions. Purified dsRNA, at a concentration of 300-1000 ng/μL, was injected into L4 or young adult hermaphrodites using a Narishige injection apparatus, a Parker Instruments Picospritzer II, and a Nikon Eclipse TE300 microscope with DIC optics. Excess dsRNA was aliquoted to avoid repeated freeze-thaw cycles and stored at –80°C. For complete knockdown, injected worms were allowed to recover on a seeded NGM plate for at least 28 hours at room temperature before harvesting embryos for imaging. For partial RNAi, worms were collected for imaging after 6-8 hours.

### Image acquisition

Embryos were dissected from gravid adults and mounted on poly-L-lysine-coated coverslips in *C. elegans* egg buffer, then sealed with VALAP (a 1:1:1 mixture of vaseline, lanolin, and paraffin). For ventral mounts, glass beads (∼23 µm in diameter, Whitehouse Scientific Monodisperse Standards, MS0023) and clay feet were used to create separation between the slide and coverslip. For lateral mounts, 2.5% agar pads were used. Fluorescence imaging was carried out using a Hamamatsu ORCA QUEST qCMOS camera mounted on a Nikon Eclipse Ti inverted confocal microscope equipped with a Yokogawa CSU-X1 spinning-disk scan head. Images were captured at room temperature using a 60× 1.4 NA oil immersion lens and lasers at 445, 488, 514, and 561 nm wavelengths. Some images were taken with a 1.5× magnifier. Z-stacks with a step size of 0.5 to 1 μm were collected every 2-3 minutes. Four-dimensional (4D) DIC video microscopy was performed on a Nikon Eclipse 800 microscope using a Diagnostic Instruments SPOT2 camera, capturing images with a 60× 1.4 NA oil objective every minute at 1-μm optical sections. Images were analyzed using MetaMorph software (MetaMorph, Inc.).

### Quantification of protein localization at subcellular regions

Signal intensity at different subcellular regions was collected in ImageJ/FIJI (Schindelin et al. 2012). For contacts between cells, the freehand line tool was used with a line width of 5 pixels (∼0.44 μm) to select the region of interest and measure the mean gray value. The Ep-P_4_ contact refers to the region where Ep and P_4_ cells touch. The other cell contacts measurement reflects contacts between Ea and Ep with each other and their neighboring cells excluding P_4_. For cytoplasmic signal intensity in Ea and Ep, a region of interest within each cell was drawn, excluding the cell boundaries and nucleus. Background subtraction was performed post hoc, by manually subtracting the mean gray value of a region outside of the embryo, to account for camera noise. Measurements were collected at 0, 6, and 12 minutes following division of the neighboring mesoderm precursor cells (MSx). All measurements were normalized to the average cytoplasmic signal.

### Colocalization analysis

The extent of colocalization between two proteins was determined using ImageJ/FIJI (Schindelin et al. 2012) and the Coloc 2 plugin (https://imagej.net/plugins/coloc-2). First, images were processed with the rolling ball background subtraction algorithm (radius: 50 pixels) and a Gaussian blur filter (radius: 2 pixels) to account for noise. Regions of interest were selected as described in the previous section, however, a thicker line width of 20 pixels (∼1.78 μm) was used to ensure that we captured areas of both high and low signal intensity for comparison. The Coloc 2 plugin was performed using the Costes threshold regression method, and the Pearson correlation coefficient was collected. Measurements were collected at 6 minutes following the division of the neighboring mesoderm precursor cells (MSx).

### Quantification of uniformity of protein localization across cytoplasm

To quantify the uniformity of protein localization across the cytoplasm, we collected line scans in ImageJ/FIJI (Schindelin et al. 2012). First, images were processed with a Gaussian blur filter (radius: 2 pixels) to account for noise. Straight lines with a line width of 10 pixels (∼0.89 μm) were drawn across the cytoplasm, excluding cell-cell boundaries and the nucleus. Maximum and minimum gray values within the region of interest were measured, and the differences between them were calculated and plotted. Representative line scan plots were also provided. Measurements were collected at 6 minutes following the division of the neighboring mesoderm precursor cells (MSx).

### Quantification of RNAi efficiency

To quantitatively evaluate the efficacy of the RNAi treatments, we mounted stage-matched wild-type, knock-in, and RNAi-treated embryos side-by-side. Relative intensity measurements were obtained from the sum intensity Z-projections of the embryos in ImageJ/FIJI (Schindelin et al. 2012). Intensity was measured by drawing a region of interest around the embryo and measuring the mean gray value, then manually subtracting the mean gray value of a background region to account for camera noise.

### Quantification of change in protein localization upon on RNAi treatment

For all RNAi-related experiments, control and RNAi-treated embryos were mounted side by side to facilitate quantification. To quantify changes in signal intensity due to RNAi treatment, measurements were taken in the previously described regions of interest using a line width of 5 pixels (∼0.44 μm) in ImageJ/FIJI (Schindelin et al. 2012). Background subtraction was performed manually by subtracting the mean gray value of a region outside the embryo to account for camera noise. Measurements were collected 6 minutes after the division of neighboring mesoderm precursor cells (MSx). Measurements at each region were normalized to the average signal intensity of the control cytoplasm.

### Establishing ectopic cell-cell contacts

The eggshell and vitelline envelope of the embryos were removed as described previously (Edgar and Goldstein 2012). Briefly, a solution containing chitinase (5 units/mL, Sigma C6137) and chymotrypsin (10 mg/mL, Sigma C4129) was used to digest the eggshell. The vitelline envelope was then mechanically stripped off. Embryos, now free of the eggshell and vitelline envelope, were generally pushed together to facilitate the formation of ectopic cell-cell contacts. These newly formed chimeras were mounted on glass coverslips with glass beads and clay feet in Shelton’s Medium and then sealed with VALAP (a 1:1:1 mixture of vaseline, lanolin, and paraffin) for imaging.

### Quantification of protein localization at ectopic contacts, endogenous contacts and contact-free cell apexes

To compare the amount of protein localized to ectopic contacts, endogenous contacts, and contact-free cell apexes, measurements were taken in these regions using a line width of 5 pixels (∼0.44 μm) in ImageJ/FIJI (Schindelin et al. 2012). Endogenous contacts and contact-free cell apexes were selected near the ectopic contacts to minimize potential variation from uneven illumination. For chimeras with multiple ectopic contacts, an average measurement for each region within the same embryo was calculated and used for statistical analysis. Background subtraction was performed manually by subtracting the mean gray value of a region outside the embryo to account for camera noise. Measurements at each region were normalized to the average signal intensity of the contact-free cell apexes.

### Statistical analysis

Comparative analyses of Gad phenotypes across different groups were conducted using Fisher’s exact test. For comparisons between two groups, unpaired t-tests were used if variances were equal, and unpaired t-tests with Welch’s correction were employed if variances were unequal. For comparisons among more than two groups, one-way ANOVA was used when variances were equal. Post-hoc Tukey’s tests were performed to identify pairwise differences when significant differences were found. If variances were unequal, Welch’s ANOVA was applied, followed by post-hoc Dunnett’s tests for pairwise comparisons in the presence of significant differences. Pearson correlation coefficients among more than two groups were compared using the Kruskal-Wallis test, and post-hoc Dunn’s tests were conducted for pairwise comparisons when significant differences were found. P-values and sample sizes (n) for each experiment are reported in the figures, figure legends, or text. All statistical analyses were performed using GraphPad Prism 10.

## Data availability statement

The data are available from the corresponding author upon reasonable request.

## Acknowledgments

We thank members of the Goldstein lab, Dan Dickinson, Ed Munro, Dave Reiner, Martha Soto, and Jessica Sullivan-Brown for helpful feedback, discussion, and/or critical reading of the manuscript. This work was supported by a Maximizing Investigators’ Research Award (R35GM134838 to B.G.) from the National Institutes of Health. T.N.M.-K. was supported by postdoctoral fellowships from the American Cancer Society (A24-0591-001) and the L’Oréal For Women in Science Program. E.A.B was supported by a Summer Undergraduate Research Fellowship from the UNC Chapel Hill Office for Undergraduate Research. Some strains were provided by the CGC, which is funded by NIH Office of Research Infrastructure Programs (P40 OD010440).

## Figure legends

**Figure S1.**
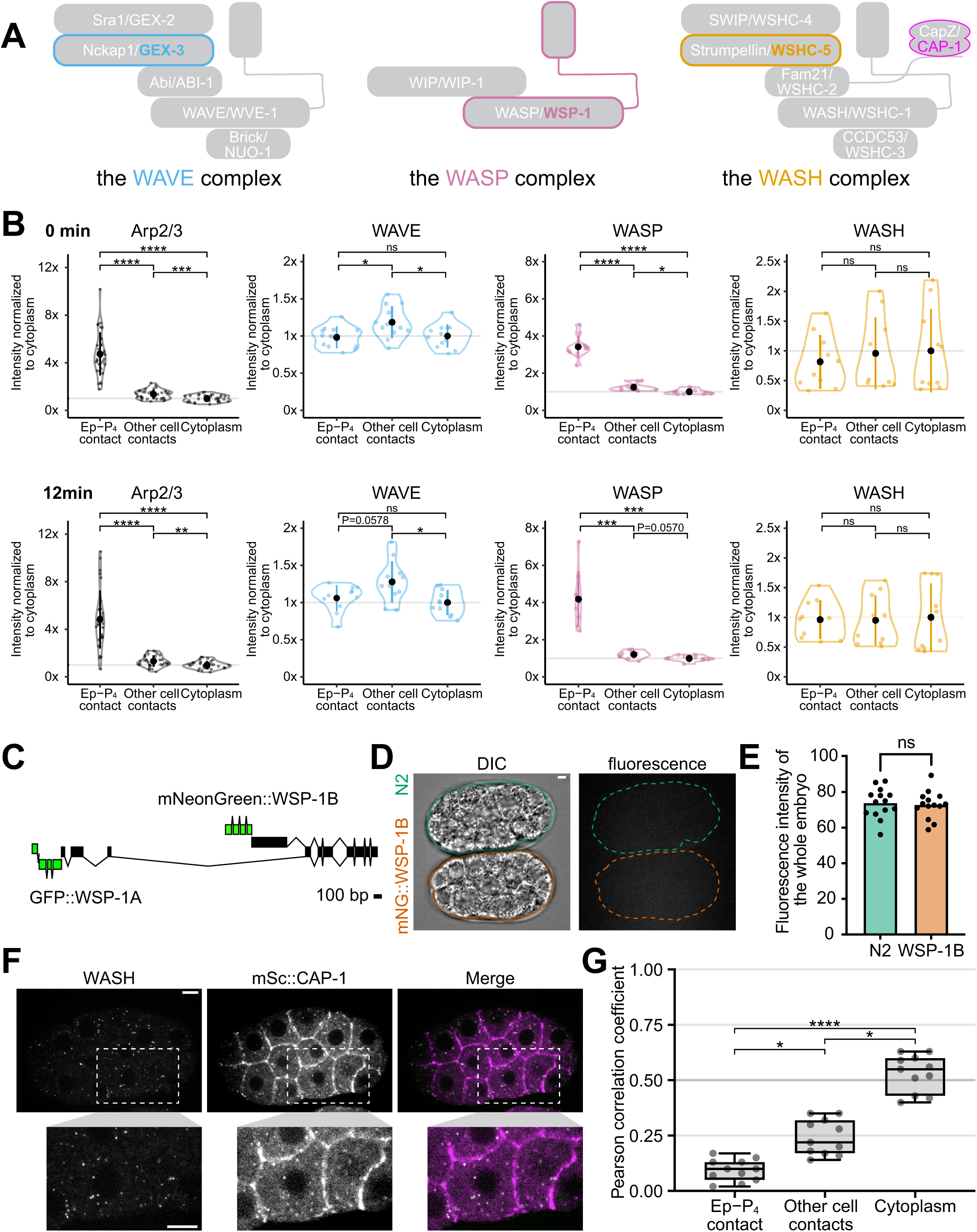
Characterization of the NPFs during apical constriction. (A) Schematic of the WAVE, WASP, and WASH complexes with endogenously-tagged components highlighted with colored outlines. (B) Violin plots depicting normalized fluorescence intensity of Arp2/3, WAVE, WASP, and WASH at Ep/P_4_ contacts, cell-cell contacts, and the cytoplasm. Insets of Arp2/3 and WASP highlight the difference between the signal at the cell-cell contacts and the cytoplasm. Measurements were collected 0 and 12 minutes after the division of neighboring mesoderm precursor cells (MSx). (center dot, mean; vertical line, s.d.; outline, the distribution of the data; n ≥ 10 embryos) (C) Schematic of the endogenous wsp-1 locus fused to GFP or mNeonGreen at its N-termini of the isoforms encoding WSP-1A or WSP-1B. (D) DIC (left) and fluorescence (right) micrographs of N2 control and mNeonGreen::WSP-1B embryos from a lateral view. (E) Bar plot depicting relative intensity measurements of whole embryos from N2 and mNeonGreen::WSP-1B worms (n = 15 embryos). (F) Co-visualization of WASH and mScarlet-I::CAP-1 from a lateral view. Areas within the white boxes in the upper panel are enlarged in the lower panel to better visualize colocalization between the two proteins. (G) Quantification of colocalization, with a box plot reporting Pearson correlation coefficients at Ep/P_4_ contacts, cell-cell contacts, and the cytoplasm. (center line, median; box, IQR; whiskers, min/max range; n = 10 embryos). P-values reported in Figures B were calculated using either Tukey’s test or Dunnett’s test, depending on whether the variances between groups were equal. P values reported in Figure E were calculated with an unpaired t-test. P values reported in Figures G were calculated with a Dunn’s test.(*p<0.05, **p<0.01, ***p<0.001, ****p<0.0001) Scale bar: 5 µm.

**Figure S2.**
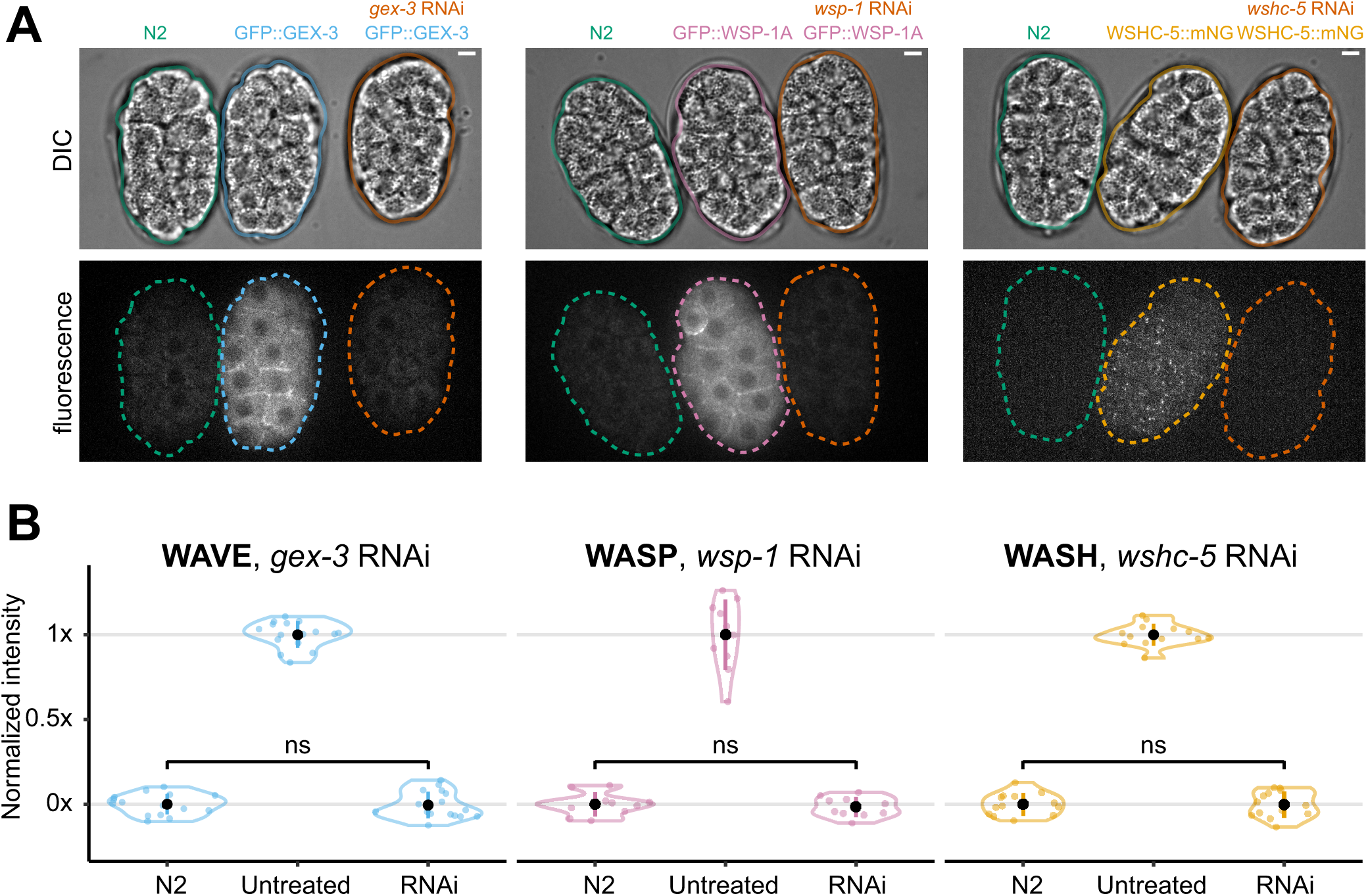
Quantification of fluorescence in wild-type/knock-in/RNAi-treated embryos permits verification of knockdown effectiveness. (A) DIC (upper) and fluorescence (lower) micrographs of stage-matched wild-type, knock-in, and RNAi-treated embryos mounted side-by-side from a lateral view. Scale bar: 5 µm. (B) Violin plots depicting normalized relative intensity measurements of whole embryos from wild-type, knock-in, and RNAi-treated embryos. Average fluorescence intensity in wild-type embryos is set to 0%, and in knock-in embryos is set to 100%. P values were calculated using a Dunnett’s test. (center dot, mean; vertical line, s.d.; outline, the distribution of the data; n ≥ 10 embryos)

**Figure S3.**
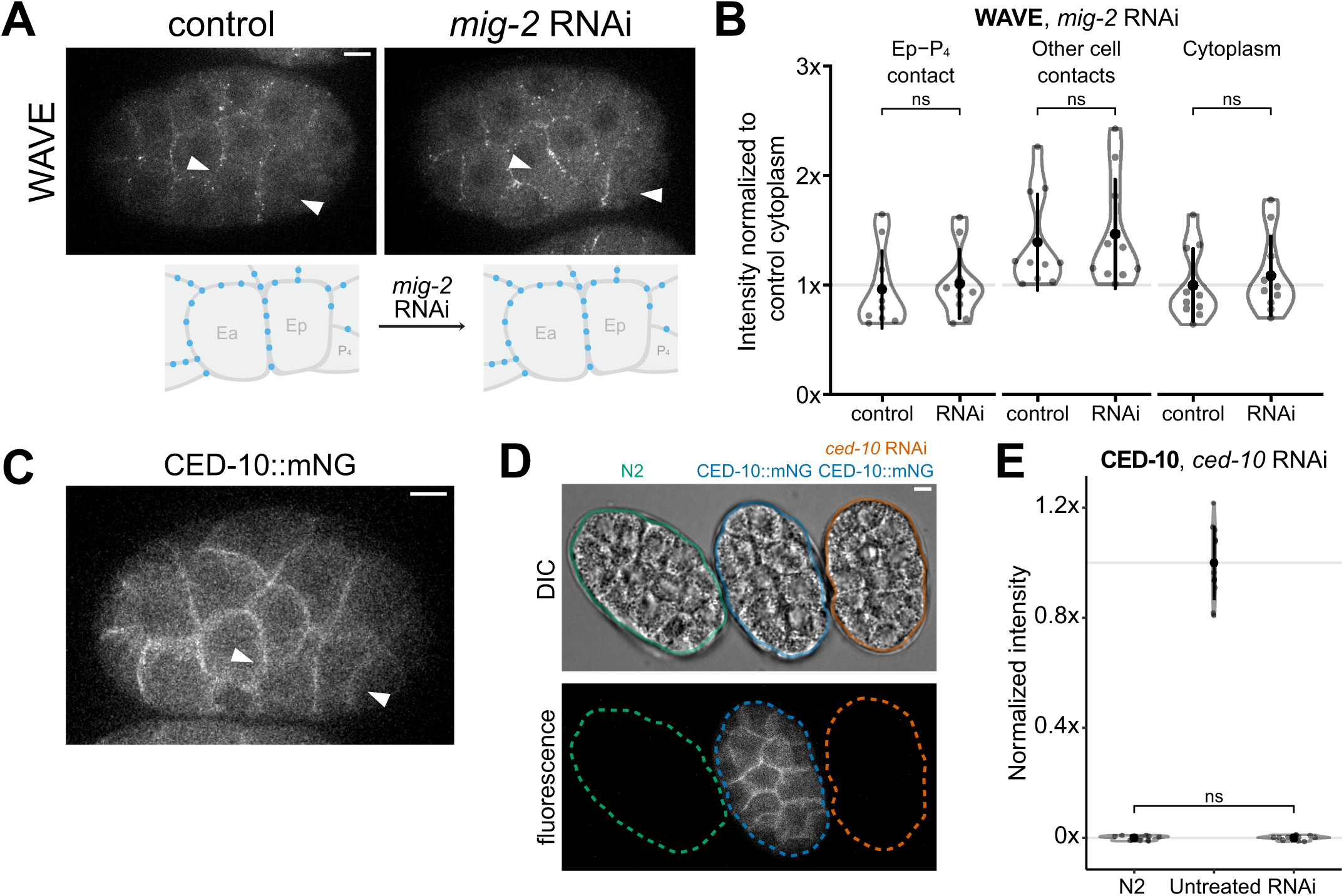
Characterization of the Rac proteins during apical constriction. (A) Micrographs from time-lapse movies depicting localization of WAVE and Arp2/3 in control and mig-2 RNAi-treated embryos from a lateral view. The diagrams underneath each micrograph highlight the observed Arp2/3 localization in E and neighboring cells. (B) Violin plots reporting changes in WAVE and Arp2/3 localization at Ep/P_4_ contacts, cell-cell contacts, and the cytoplasm upon RNAi depletion of MIG-2. (center dot, mean; vertical line, s.d.; outline, the distribution of the data; n = 10 embryos) (C) Micrographs from time-lapse movies depicting localization of mNeonGreen::CED-10 from a lateral view. White arrowheads point to Ea and Ep cells. (D) DIC (up) and fluorescence (down) micrographs of stage-matched wild-type, knock-in, and ced-10 RNAi-treated embryos mounted side-by-side from a lateral view. (E) Violin plot depicting normalized relative intensity measurements of whole embryos from wild-type, knock-in, and ced-10 RNAi-treated embryos, with average fluorescence intensity in wild-type embryos set to 0% and knock-in embryos set to 100%. (center dot, mean; vertical line, s.d.; outline, the distribution of the data; n = 11 embryos) P values reported in Figure B were calculated with an unpaired t-test, and P values reported in Figure E were calculated with a Dunnett’s test. (*p<0.05, **p<0.01, ***p<0.001, ****p<0.0001) Scale bar: 5 µm.

**Figure S4.**
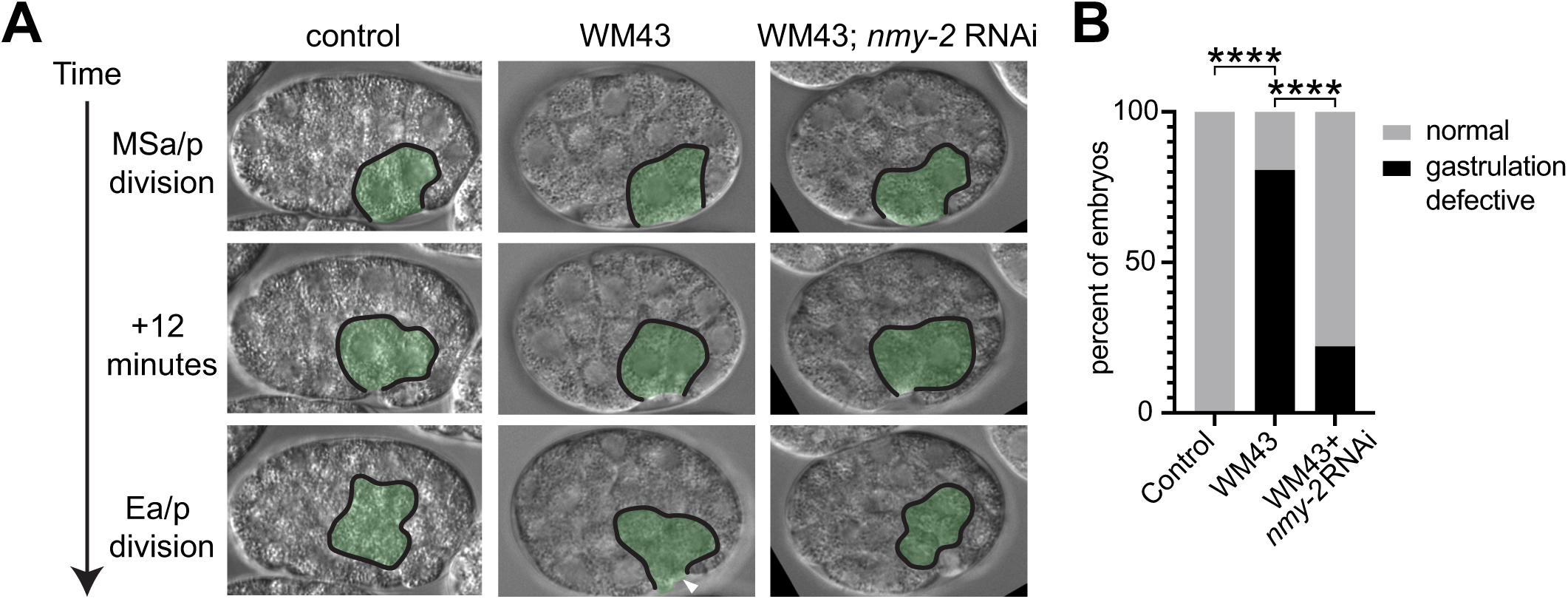
Myosin II partial RNAi rescued gastrulation defects caused by WAVE depletion. (A) Micrographs from time-lapse DIC movies in three backgrounds with time on the left from MSa/p cell division. E lineage cells are outlined and pseudocolored in green. Gastrulation defects (E cells dividing before being fully covered by neighboring cells) are indicated with white arrowheads. An enclosed outline and absence of the arrowhead indicate that endodermal precursors became internalized at the 2E stage, as in wild-type embryos. (B) The bar graph summarizes the percentage of embryos that have gastrulation defects. P values were calculated with Fisher’s exact test. (*p<0.05, **p<0.01, ***p<0.001, ****p<0.0001)

## Supplemental material

**Table S1:**
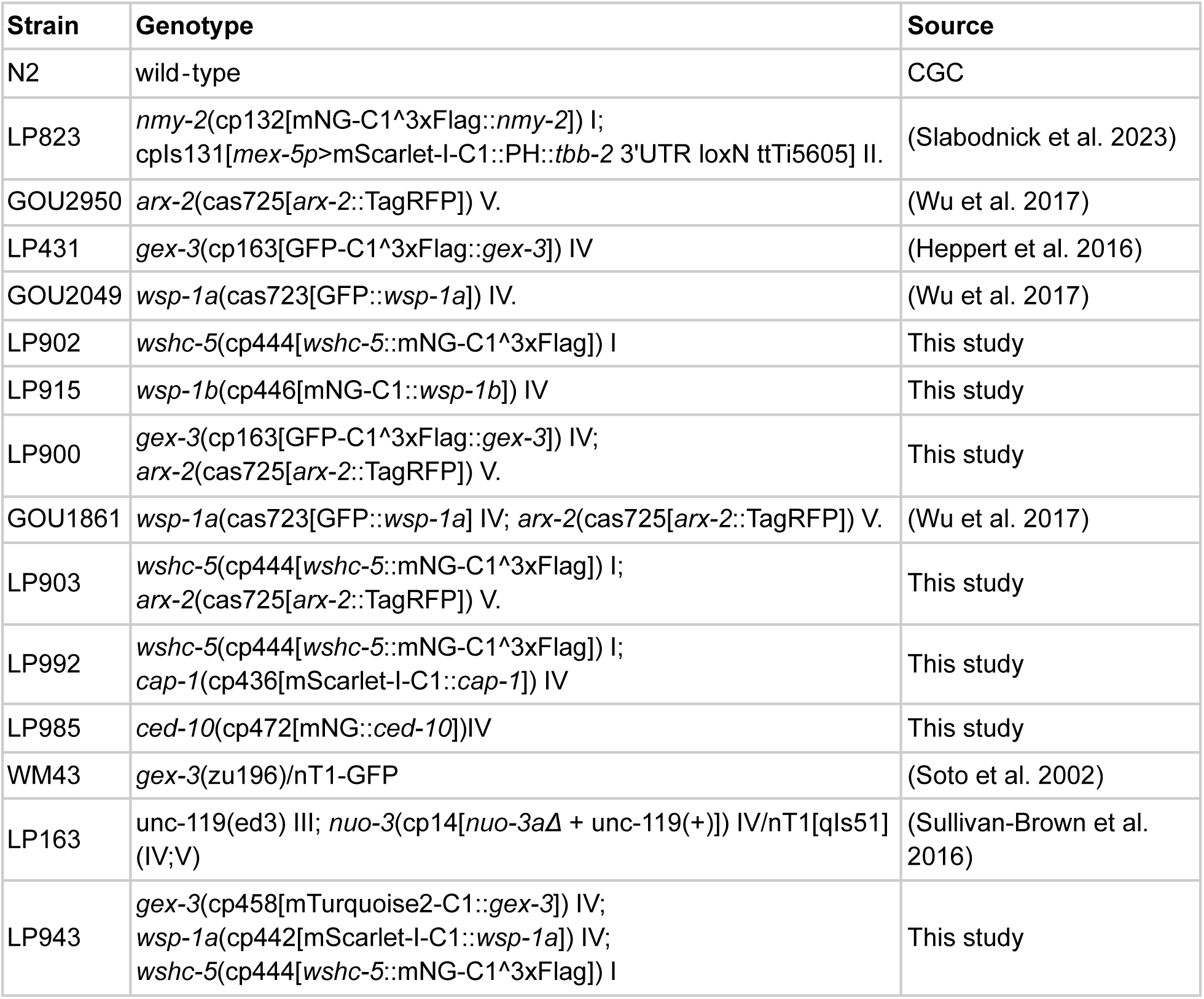
Strains. Names, genotypes, and associated references for the strains used in this study.

**Table S2:** Embryo hatching rates. Hatching rates for strains used in this study. N refers to the number of embryos examined.

| Strain | Genotype | Hatching rate | N |
| --- | --- | --- | --- |
| GOU2950 | cas725[arx-2::TagRFP] V. | 100% | 129 |
| LP431 | gex-3(cp163[GFP-C1<sup>3</sup>xFlag::gex-3]) IV | 99% | 111 |
| GOU2049 | wsp-1a(cas723[gfp::wsp-1a]) IV. | 100% | 109 |
| LP902 | *wshc-5*(cp444[*wshc-5*::mNG-C1<sup>3</sup>xFlag]) I | 100% | 110 |
| LP915 | wsp-1b(cp446[mNG-C1::wsp-1b]) IV | 100% | 175 |
| LP985 | ced-10(cp472[mNG::CED-10])IV | 91% | 202 |
| LP943 | gex-3(cp458[mTurquoise2-C1::gex-3]) IV;<br>wsp-1(cp442[mScarlet-I-C1::wsp-1]) IV;<br>*wshc-5*(cp444[*wshc-5*::mNG-C1<sup>3</sup>xFlag]) I | 100% | 117 |

**Table S3:**
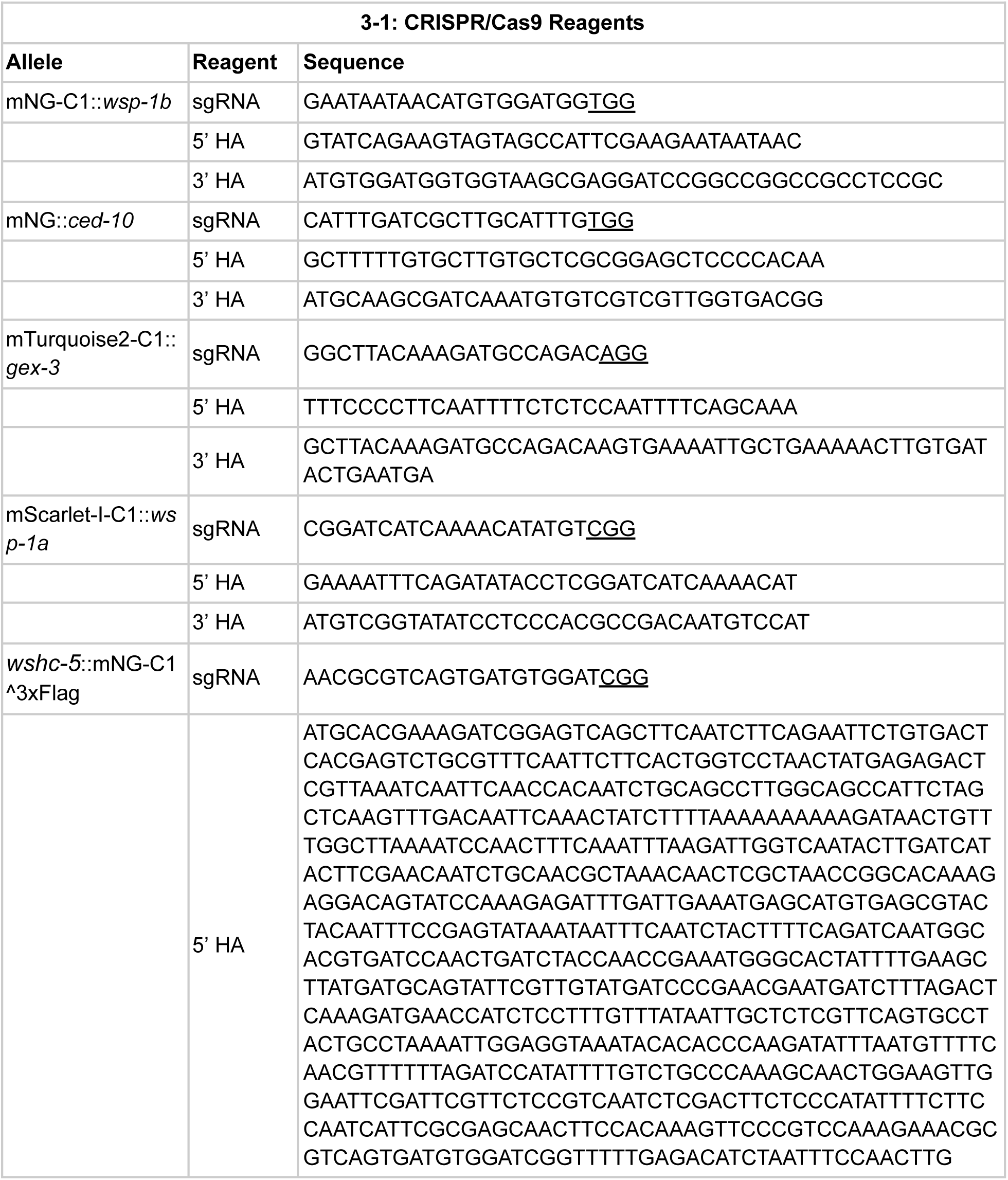
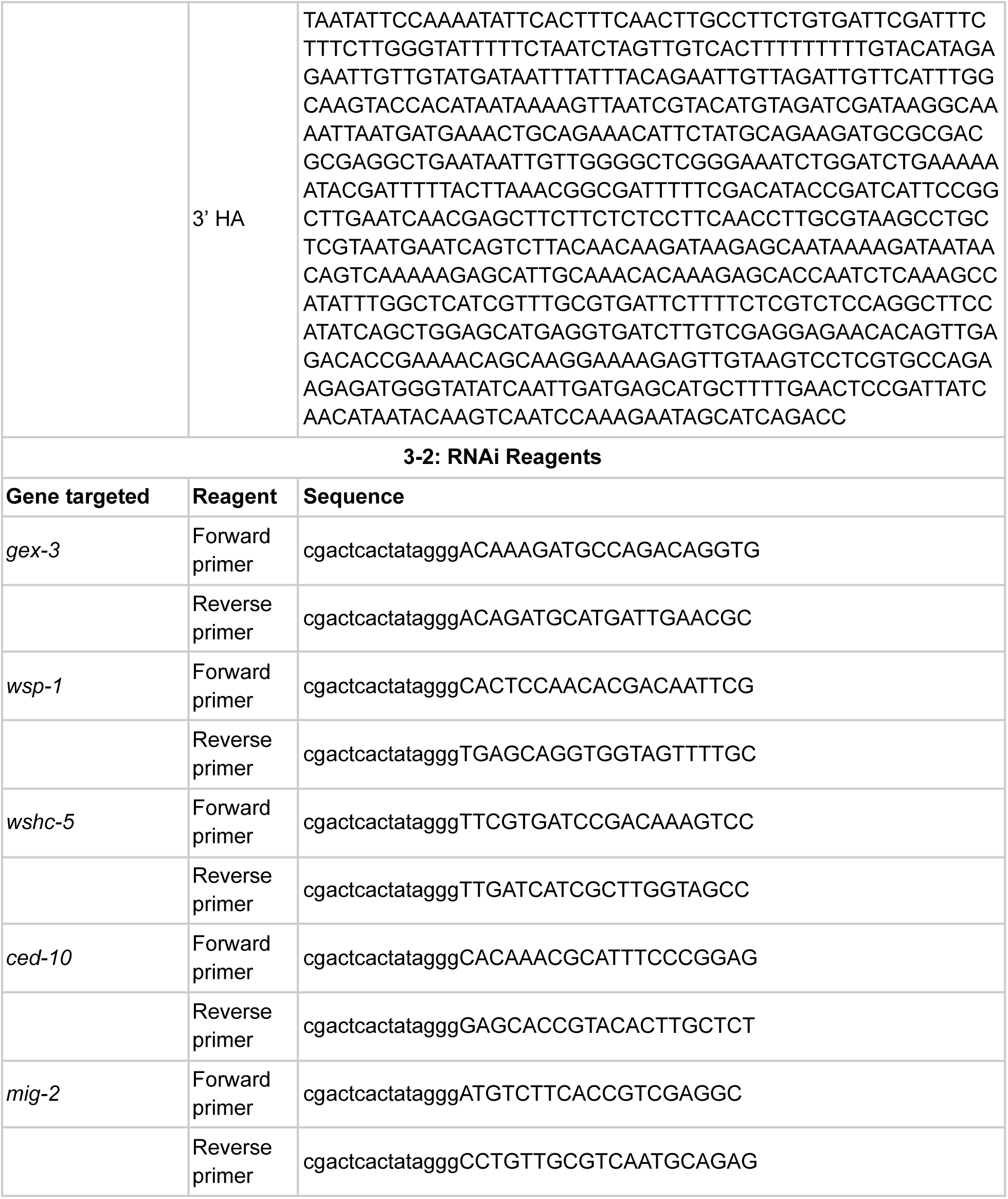
Sequences of CRISPR/Cas9 and RNAi reagents. S3-1, sequences of the homology arms (HA) and single guide RNAs (sgRNA) with PAM sequences underlined. S3-2, sequences of primers designed to amplify approximately 1000 bp of the protein-coding sequence of each target gene.

**Table S4:** Time between MS and E division in all conditions examined. None of the RNAi treatments lead to a significant change. P-values were calculated using a Tukey test.

| Strain | RNAi | Embryo number | Time between MS and Ea division (min) | P value | Time between MS and Ep division (min) | P value |
| --- | --- | --- | --- | --- | --- | --- |
| N2 | untreated control | 27 | 20.26 ± 1.23 | NA | 22.26 ± 1.20 | NA |
| N2 | WAVE | 46 | 19.96 ± 1.10 | ns | 21.78 ± 1.25 | ns |
| N2 | WASP | 41 | 20.37 ± 1.62 | ns | 21.98 ± 1.97 | ns |
| N2 | WASH | 28 | 21.04 ± 1.07 | ns | 22.71 ± 1.12 | ns |
| N2 | WAVE + WASP | 64 | 20.34 ± 1.44 | ns | 22.36 ± 1.33 | ns |
| N2 | WAVE + WASH | 48 | 21.04 ± 1.40 | ns | 22.88 ± 1.06 | ns |
| N2 | WASP + WASH | 35 | 20.29 ± 1.74 | ns | 22.40 ± 1.68 | ns |
| N2 | All NPFs RNAi | 44 | 20.75 ± 1.42 | ns | 22.75 ± 1.38 | ns |
| N2 | CED-10 | 39 | 20.44 ± 1.67 | ns | 22.36 ± 1.63 | ns |
| WM43 | untreated control | 26 | 19.54 ± 1.10 | ns | 21.35 ± 1.06 | ns |
| WM43 | NMY-2 | 27 | 19.81 ± 1.04 | ns | 21.48 ± 1.48 | ns |

## Notes

### Competing Interest Statement

The authors have declared no competing interest.

